# The Cia1 and Cia2 subunits of the CTC mediate recognition of apo-FeS proteins with a C-terminal targeting complex recognition motif

**DOI:** 10.1101/2025.03.25.645274

**Authors:** A Buzuk, MD Marquez, JV Ho, Y Liu, B Wang, CC Qi, DL Perlstein

## Abstract

The cytosolic iron-sulfur cluster assembly (CIA) targeting complex is responsible for maturation of cytosolic and nuclear iron-sulfur enzymes, numbering >30 proteins critical for fundamental processes such as DNA replication and repair. Up to 25% of these client proteins terminate in a targeting complex recognition (TCR) motif. This carboxy-terminal tripeptide motif recruits the CIA targeting complex (CTC) to the client so that the metallocluster can be inserted. Herein, we use a combination of computational, biochemical and biophysical approaches to determine that the clients bearing a TCR motif docks at the interface of the Cia1 and Cia2 subunits of the CTC. Thus, mutations destabilizing the Cia1-Cia2 complex also disrupt TCR-based client identification by the CTC. Our study also reveals that the understudied human Cia2 paralog CIAO2A, which is proposed to be a specific targeting factor for iron regulatory protein 1, can recruit clients terminating in the TCR peptide. These data signal that CIAO2A plays a more general role in iron-sulfur protein maturation than previously appreciated. Taken together, our findings deepen our understanding of the molecular basis for client recognition by the CTC that is critical to understand the impact of CIA function in human health and disease.

## Introduction

Iron-sulfur (Fe-S) proteins are required for countless essential processes in which their metalloclusters perform a myriad of functions, such as electron transfer during photosynthesis and oxidative phosphorylation; catalysis in nitrogen fixation; and non-redox functions such as a sensor for iron homeostasis.^1^ The *de novo* biogenesis and delivery of these metallocofactors requires dedicated biochemical pathways.^2-6^ Although several Fe-S biosynthesis machineries have been identified, they all appear to follow a common mechanism in which nascent [2Fe-2S] clusters are assembled from iron and sulfide.^7-9^ These [2Fe-2S] clusters can be further combined and modified to form [4Fe-4S] or more complex clusters before their delivery to an apo-protein client by a network of trafficking and targeting factors.^2-6, 10^ Unlike most metallocofactor biogenesis systems in which each metalloprotein has its own dedicated assembly factor, the Fe-S cluster targeting factors must recognize many different clients, raising the question of how specificity is encoded.

For example, the cytosolic iron-sulfur cluster assembly (CIA) system in eukaryotic organisms is responsible for the maturation of cytosolic and nuclear Fe-S proteins, including those critical for DNA replication and repair, iron homeostasis, and the biosynthesis of nucleotides and amino acids.^2^ In the late stages of CIA, the CIA targeting complex (CTC) is proposed to accept a nascent cluster from CIAO3/Nar1 (HUMAN/yeast nomenclature), the putative [4Fe-4S] cluster carrier.^11, 12^ Next, the CTC, comprising the MMS19/Met18, CIAO1/Cia1, and CIAO2B/Cia2 subunits, identifies apo-clients via direct or adaptor-mediated interactions and delivers the cluster to the CTC-bound client **(Figure 1A**).^13-15^ Several pioneering studies probing CIA client recognition indicated that each of the CTC subunits directly interacts with clients.^16-20^ However, recent work has suggested that the primary role of CIAO2B/Cia2 is to bind the Fe-S cluster that is delivered to clients, whereas MMS19/Met18 and CIAO1/Cia1 mediate client identification.^11, 21-24^ Because the biochemical characterization of CIA factor function has lagged behind their identification via genetic approaches, we have made little progress deciphering the molecular signals guiding CIA client identification in the 14 years since discovery of the CTC.^13-15^

**Figure 1.**
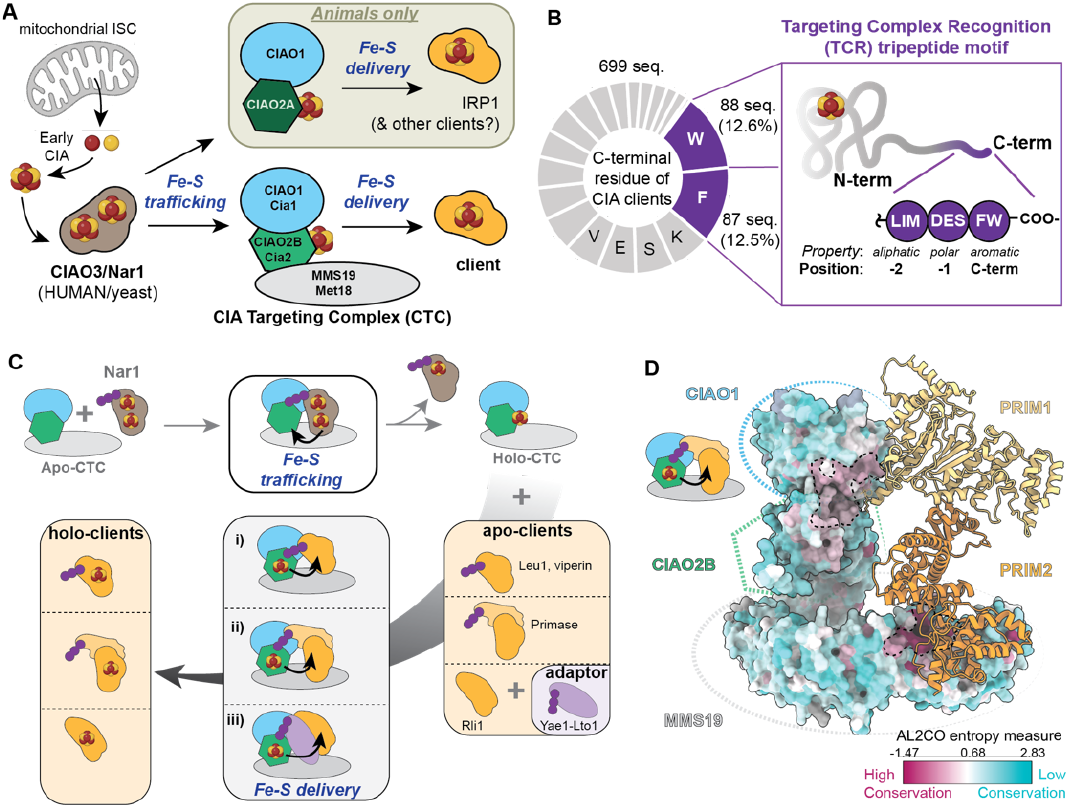
Overview of the CIA pathway and role of the TCR motif. (**A**) The CIA trafficking factor CIAO3/Nar1 (HUMAN/yeast nomenclature) donates an Fe-S cluster to CTC, which subsequently identifies apo-clients and inserts their metallocofactor. The Cia2 paralog CIAO2A, which is unique to animals, is proposed to be a specific targeting factor for IRP1. (**B**) Up to 25% of CIA clients and CTC-binding proteins (n=699 sequences) end in a Trp or Phe residue, which is characteristic of the TCR motif (purple). (**C**) In our current working model of the CIA pathway, CIAO3/Nar1 binds to the CTC via its TCR motif (top), transfers an Fe-S cluster and dissociates. Next, the CTC identifies clients containing the TCR peptide (purple) via one of three modalities: the Fe-S protein itself (**i**), or its binding partner (**ii**), ends in a TCR motif, or an adaptor harboring a TCR motif transiently interacts with the apo-client (**iii**). Examples of clients recognized by each mechanism are indicated. (**D**) The primase-CTC structure showing how the conserved surfaces (dashed lines, dark pink) of CIAO1/Cia1 (top) and MMS19/Met18 (bottom) are adjacent to the PRIM1 and Fe-S binding PRIM2 subunits of primase, respectively.^24, 25^

The first insight into this important question was provided by discovery of the targeting complex recognition (TCR) motif.^25^ This tripeptide with the [LIM]-[DES]-[FW] consensus sequence is found at the C-terminus of 20-25% of CIA clients (**Figure 1B**). When present, the TCR motif is sufficient for recruitment of the CTC so that the metallocluster can be inserted (**Figure 1C**).^25^ Clues as to where the TCR peptide docks on the CTC were provided by the structure of the CTC-primase (PRIM1-PRIM2) complex, showing the PRIM2 subunit of primase is adjacent to MMS19/Met18 and that the TCR-peptide bearing PRIM1 subunit was closest to the third β-propeller domain of CIAO1/Cia1 (**Figure 1D**).^24^ The hypothesis that Cia1 contacts to the TCR peptide is corroborated by previous studies which reported that the CIA client Viperin (RSAD2) binds Cia1 via Viperin’s C-terminal tryptophan.^26^ Taken together, these studies provided evidence for Cia1’s role recruiting clients ending in the TCR tripeptide, but they left Cia2’s function largely unresolved.

Herein, we use a combination of biochemical and computational studies to determine that the TCR motif binds to the CTC at the interface of the CIAO1/Cia1 and CIAO2/Cia2 subunits. Two key arginine residues, one from Cia1 and one from Cia2, are identified as hotspot residues, each contributing more than 1.4 kcal/mol in binding energy. Since these residues are absolutely conserved in animals, fungi, and plants, our study establishes a universal mechanism of CIA client identification. Furthermore, these TCR peptide binding residues are conserved in the paralogous pair of Cia2 proteins found in animals, called CIAO2A and CIAO2B. CIAO2B is the proposed housekeeping subunit, serving as a general targeting factor for CIA clients, whereas CIAO2A is proposed to function as a specific targeting factor for IRP1 (**Figure 1A**), a cytosolic Fe-S protein that regulates iron homeostasis.^18^ Since human CIAO1-CIAO2B is able to perform TCR peptide recognition *in vitro*, our findings suggest that CIAO2A facilitates maturation of a wider variety of metazoan Fe-S proteins than previously hypothesized.

## RESULTS

### Cia1 is required to bind the TCR-tripeptide

We were intrigued that CIOA3/Nar1 and a subset of CIA clients terminate with a TCR motif because it suggested that the protein proposed to donate the Fe-S cluster to the CTC (Nar1) and apo-clients which receive an Fe-S cluster from the CTC, such as Leu1, bind to the same site on the CTC. To test if *Saccharomyces cerevisiae* (*Sc*) Nar1 and *Sc*Leu1 compete for binding to the CTC, the *Sc*CTC-Leu1 complex was incubated with increasing amounts of *Sc*Nar1 and the mixture was passed through streptactin resin, which binds the strep-tagged Cia2 bait. As the amount of Nar1 in the eluate increased, we observed a corresponding decrease in the amount of Leu1 (**Figure 2A** and **S1A**), indicating that the interaction of Nar1 and Leu1 with the CTC is mutually exclusive.

**Figure 2.**
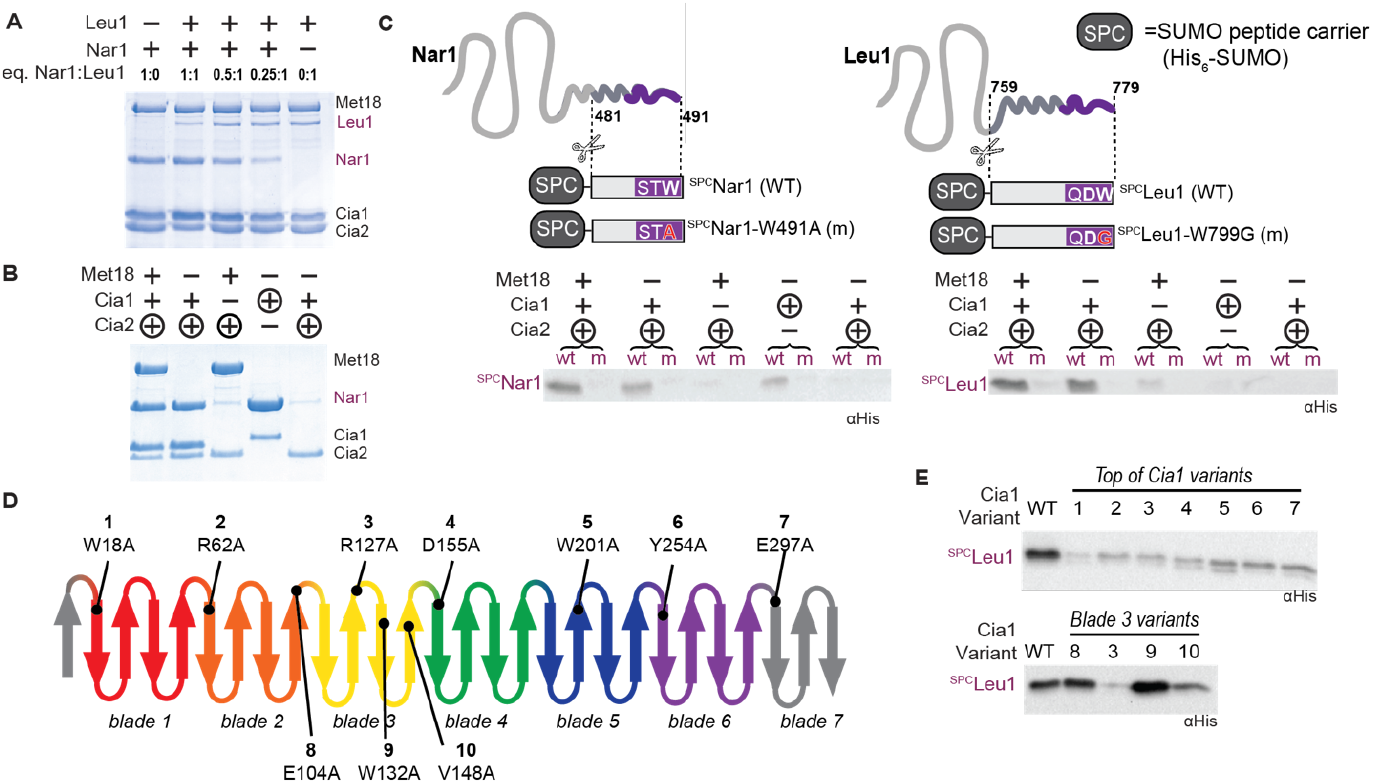
Cia1 is required to bind the TCR peptide. (**A**) Affinity copurification demonstrating that *Sc*Nar1 competes with *Sc*Leu1 for binding to the *Sc*CTC. Cia1, Cia2 (strep-tagged bait), Met18, and Leu1 were mixed with Nar1 (0.25-1.0 molar equivalents (eq.) relative to Leu1). The streptactin eluate was analyzed via SDS-PAGE. (**B**) The indicated *Sc*CTC subunit(s) (strep-tagged bait is indicated with a circle) were incubated with *Sc*Nar1. SDS-PAGE analysis of the streptactin resin eluate shows that the Cia1 subunit is sufficient to bind Nar1. (**C**) Copurification experiments as in **B**, except the indicated SPC-variants were used in place of Nar1 and eluate was analyzed via western blot, detecting the His-tagged SPC. Controls in which the C-terminal Trp was mutated (m) to Ala (^SPC^Nar1-W491A) or Gly (^SPC^Leu1-W779G) ensured any interactions detected depend on the TCR peptide. (**D**) A schematic of Cia1’s seven β-propeller domains. The location of each *Sc*Cia1 variant along the top face (1-7) or the side of blade 3 (8-10) are indicated. (**E**) The indicated *Sc*Cia1 variant was mixed with ^SPC^Leu1 and *Sc*Cia2 (strep-tagged bait) and copurification experiments carried out as in **C**.

Next, we substituted *Sc*Nar1’s C-terminal Trp with Ala and repeated the copurification experiment with the CTC. Since less Nar1 TCR-peptide variant was found in the eluate (**Figure S1B**), we concluded Nar1’s C-terminal tail contributes to recruitment of the CTC. To pinpoint which subunit(s) of the CTC bind to Nar1, we performed copurification experiments using the CTC subunits alone or in combination. We observed Nar1 eluting with Cia1, Cia1-Cia2, and the full CTC (**Figure 2B** and **S1C**). From these experiments, we concluded Nar1’s binding to the CTC is centered on the Cia1 subunit.

To ascertain whether or not the TCR peptide is sufficient for the interaction with Cia1, SUMO peptide carrier (SPC) constructs of Nar1 and Leu1 were used as prey proteins in the copurification assay.^25^ These SPC proteins have the last 10 residues of Nar1 (residues 482-491) or the last 21 residues of Leu1 (759-779) appended to the C-terminus of SUMO (**Figure 2C**). Previous work demonstrated these SPC-TCR peptide fusion proteins bind the CTC *in vitro*.^25^ Since these SPC constructs are small (20 kDa) compared to full-length Nar1 (54 kDa) and Leu1 (86 kDa), the increased sensitivity of a western blot was needed to detect ^SPC^Nar1 or ^SPC^Leu1 in the eluate. Negative controls employing C-terminal tryptophan substituted SPC variants ensured any observed interactions depend on the TCR peptide. Copurification experiments using *Sc*CTC subunits alone or in combination revealed that both ^SPC^Leu1 and ^SPC^Nar1 eluted with the Cia1-Cia2 complex in a tryptophan-dependent manner (**Figure 2C** and **S2C-D**). ^SPC^Nar1 also eluted with Cia1 in the absence of any other CTC subunits (**Figure 2C** and **S2C**).

Since previous studies suggested Cia1’s top face or the side of its third β-propeller domain (yellow, **Figure 2D**) as potential client binding sites,^24, 27^ we used alanine scanning mutagenesis of conserved, surface exposed residues in these regions to identify the TCR peptide binding residues. Every substitution made along the top of Cia1 disrupted the interaction of *Sc*Cia1-Cia2 with ^SPC^Leu1, whereas substitutions along the side of blade 3 had little to no effect (**Figure 2E** and **S2**). Several of these Cia1 variants also affected interaction with ^SPC^Nar1 (**Figure S1B**), supporting the hypothesis that they are involved in TCR peptide binding.

Since the top-face of Cia1 binds to Cia2 and several Cia1 variants appeared to lose interaction with Cia2 in addition to the TCR peptide (**Figure S2**), we sought to confirm that changes in TCR peptide binding were not due to nonspecific effects, such as destabilizing Cia1 or the Cia1-Cia2 complex. To quantitatively interrogate Cia1-Cia2 complex formation, we employed a microscale thermophoresis (MST) binding assay (**Figure S3** and **Table S1**). To develop this assay, we determined that titration of fluorescently labeled Cia1 with Cia2 or vice versa resulted in similar, low nanomolar dissociation constants (*K*_*D*_) (**Table S1** and **Figure S3**). Next, we showed that a Cia2 variant defective in Cia1 binding (E208A)^28^ exhibited an increased *K*_*D*_ value (**Table S1**). Finally, we removed the first 102 residues of Cia2 (Δ102) to increase the Cia2 stability without impacting Cia1 affinity (**Table S1**).^28^ Consistent with our copurification analysis, the W18A and R62A Cia1 variants reach resulted in a >300-fold decreased affinity for Cia2. To our surprise, the W201A and Y254A variants also disrupted Cia2 binding, with ≥10-fold increase in the *K*_*D*_ value, despite having no impact on the Cia1-Cia2 interaction as monitored by the qualitative copurification assay (**Figure S3** and **Table S1**). Of the Cia1 variants tested, only ^R127A^Cia1 disrupted binding to ^SPC^Leu1 without impacting affinity for Cia2. Taken together, these results suggest that R127, which sits atop Cia1’s third β-propeller domain, directly contacts the TCR peptide.

Finally, we incorporated each Cia1 variant into the CTC and tested binding to full-length Nar1 or Leu1. Although Cia1 variants disrupting interaction with ^SPC^Leu1 also diminished interaction of the CTC with full-length Leu1 (**Figure S4A**), Nar1 and ^SPC^Nar1 yielded contradictory results. The R127A and D155A Cia1 variants disrupted diminished the amount of ^SPC^Nar1 in the eluate, however none of the Cia1 variants tested, including the charge swapped R127E variant, completely blocked interaction with full-length Nar1 (**Figure S1B** and **S4B**). These observed differences suggest Nar1 binds to the CTC via a bipartite interaction requiring the TCR motif and another region of Nar1. The observation that mutation of Nar1’s C-terminal residue diminishes, but does not abolish, its interaction with the CTC corroborates this hypothesis (**Figure S2C**). In contrast, Leu1 requires its C-terminal Trp to interact with the CTC *in vitro* (**Figure S4A**),^25^ thus, explaining why any substitutions that affect affinity of the Leu1’s TCR motif for the CTC disrupt the Leu1-CTC complex. Overall, our alanine scanning mutagenesis study pinpoints R127 of *Sc*Cia1’s as the key residue for interaction with the TCR peptide.

### The TCR peptide binds at the Cia1-Cia2 interface

To quantitatively monitor the TCR-CTC interaction, a fluorescence anisotropy (FA) binding assay was developed using FITC-labeled peptide probe corresponding to the last 4 residues of *Sc*Leu1 (**Figure 3A**). As *Sc*Cia1 was titrated into the solution, we were surprised that we did not observe any binding to the probe. However, we did observe a change in the FA as the *Sc*Cia1-Cia2 complex was titrated into a solution containing the FITC-probe. Although the solubility of the complex prevented the FA signal from reaching saturation, the data that was fit to determine an apparent *K*_*D*_ value of 86±0.4 µM (**Figure 3A** and **Table S2**). The ^R127A^Cia1-Cia2 variant resulted in a ≥200-fold lowered affinity for the FITC-4mer as compared to the WT complex (**Figure 3A** and **Table S2**). Although promising, the low solubility and cumbersome purification of ScCia1-Cia2 limited its utility.

**Figure 3.**
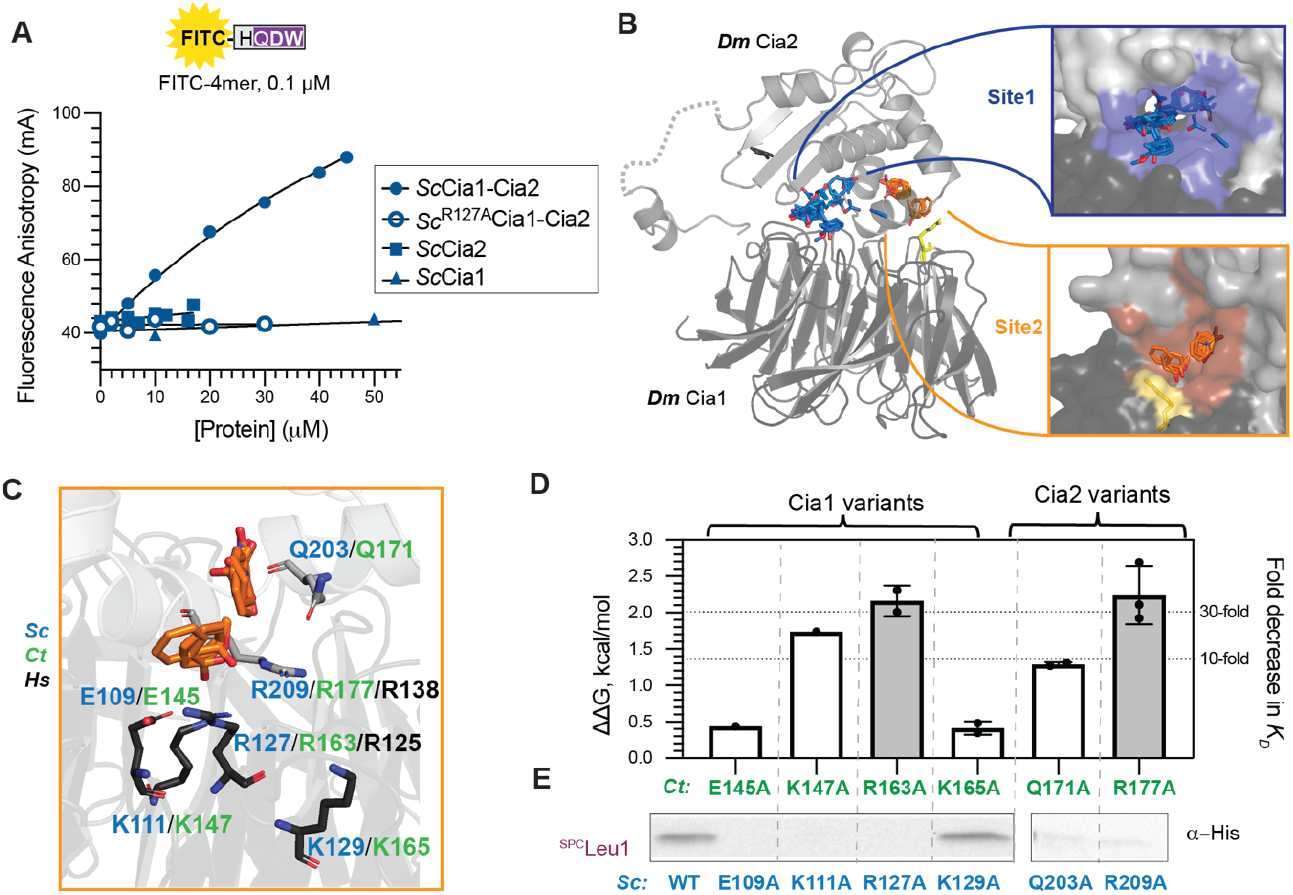
The TCR peptide binds at the Cia1-Cia2 interface. (**A**) The FITC-4mer probe (top) was titrated with the indicated *Sc* protein(s). The apparent *K*_*D*_ for the protein-peptide interaction was determined by fitting to Equation 2. Results shown are representative of at least two independent determinations. (**B**) FTMap and PrankWeb analysis of *Dm*Cia1-Cia2 (6TBN)^24^ identified two potential TCR peptide binding sites. Site 2 (orange) was adjacent to Cia1 Arg residue (yellow, corresponding to R127 of *Sc*Cia1) involved in binding the TCR peptide. (**C**) Detailed view of Site 2, including the FTMap probes (orange sticks) and surrounding residues from Cia2 (light grey sticks) and Cia1 (dark grey sticks) identified by PrankWeb. Residues mutated to Ala in *Sc, Ct* and *Hs* proteins are labeled in blue, green, and black, respectively. (**D**) The FA assay (as in **A**) was performed with the indicated *Ct* variant and the decrease in binding energy (ΔΔG) was determined. The error bars are standard deviation of at least two independent measurements. (**E**) Affinity copurification experiments (as in **Figure 2E**) testing interaction of the indicated *Sc*Cia1-Cia2 variant with ^SPC^Leu1.

To overcome these limitations, we coexpressed Cia1 and Cia2 from the thermophilic fungi *Chaetomium thermophilum* (*Ct*), purified their complex, and utilized *Ct*Cia1-Cia2 in the FA assay (**Figure S5**). Gratifyingly, the increased solubility of *Ct*Cia1-Cia2 as compared to the *Sc* orthologs allowed FA signal to reach saturation. *Ct*Cia1-Cia2 binds the FITC-4mer with an apparent *K*_*D*_ value of 25±3.2 µM. Additionally, R163 of *Ct*Cia1, analogous to R127 of *Sc*Cia1, was critical for the interaction with the FITC-probe (**Figure S5A** and **Table S2**), demonstrating that information learned with the *C. thermophilum* proteins will be applicable to understanding other Cia1-Cia2 orthologs.

Next, we applied this more robust FA assay to identify the minimal components required for the interaction of the TCR peptide with the Cia1-Cia2 complex. First, we determined the *Ct*Cia1-Cia2 binds a FITC probe derived from the last 21 amino acids of *Sc*Leu1 with comparable affinity as the FITC-4mer (**Figure S5B-C**). Thus, residues preceding the TCR motif do not strongly contribute to the binding energy. We also tested whether Met18 contributes to interaction of the CTC with the TCR peptide by monitoring the affinity of *Ct*Cia1-Cia2 for the FITC-4mer in the presence and absence of Met18. Since we were unable to isolate *Ct*Met18, we utilized *Sc*Met18. Affinity copurification experiments demonstrated that *Sc*Met18 can bind to *Ct*Cia1-Cia2, but *Sc*Met18 had no measurable impact on the affinity of *Ct*Cia1-Cia2 for the FITC-4mer (**Figure S5D-E**). Considering the conservation of the TCR motif across the eukaryotic domain of life, the small size of the tetrapeptide, and sufficiency of *Ct*Cia1-Cia2 for the binding, these experiments strongly suggested that the TCR binding pocket will be correspondingly compact and lined with conserved residues from both Cia1 and Cia2.

To identify the binding pocket for the TCR motif, two distinct computational approaches, FTMap and PrankWeb, were used.^29, 30^ In FTMap, small organic probes are docked on the protein surface to identify probe clusters, which are further grouped into consensus sites (CSs), corresponding to regions with a high probability of mediating interactions. FTMap analysis of *Drosophila melanogaster* (*Dm*) Cia1-Cia2 complex^24^ revealed two major CSs at the Cia1-Cia2 interface (**Figure 3B** and **Table S3**). Excitingly, CS2 was adjacent to the R127 of *Sc*Cia1, identified as critical for binding the TCR peptide. CS2 also comprised many aromatic probes, which could be mimicking the TCR motif’s terminal aromatic residue (**Table S3**). Encouragingly, PrankWeb identified the same sites as FTMap despite utilizing a quite different computational approach that incorporates both sequence conservation and physiochemical properties of residues lining identified cavities (**Figure 3B** and **Table S4**). The PrankWeb predicted peptide binding pocket with the highest sequence conservation score of corresponded to FTMap’s CS2 (**Table S4**), suggesting that this region contains the TCR peptide binding site.

To test this hypothesis, conserved residues lining the pocket were mutated to alanine and the resulting variants were evaluated using qualitative and quantitative assays (**Figure 3C-E**). Of the variants tested, ^R163A^*Ct*Cia1 and ^R177A^*Ct*Cia2 each resulted in the largest (≥2 kcal/mol) decrease in binding energy (**Figure 3D** and **S6A-B, Table S2**). The K147A and Q171A *Ct*Cia2 variants also diminished affinity for the FITC-4mer probe, albeit with a smaller impact. With the exception of ^K147A^*Ct*Cia1, none of the variants affected isolation of the coexpressed *Ct*Cia1-Cia2 complex (**Figure S6C**). However, ^R163A^*Ct*Cia1, ^R177A^*Ct*Cia2, and ^Q171A^*Ct*Cia2 each diminished interaction with *Sc*Leu1 and the FITC-4mer (**Figure S6D** and **Table S2**) and analogous substitutions in the *Sc* orthologs disrupted their interaction with ^SPC^Leu1 (**Figure 3E** and **S6E-G**). These data led us to conclude that the computationally predicted Site 2 is the binding site for the TRC motif.

### *Hs*CIAO1 variants which result in a fatal neuromuscular disorder disrupt its interaction with *Hs*CIAO2A and the TCR peptide

Many animals, including *Homo sapiens* (*Hs*), encode a paralogous pair of Cia2 proteins, called CIAO2A/FAM96A (Cia2a herein) and CIAO2B/FAM96B (Cia2b herein). *Hs*Cia2b has higher sequence similarity to *Sc*Cia2 (49%) than *Hs*Cia2a (41%) and it is proposed to be the general CIA targeting factor, mediating maturation of most CIA clients.^18^ Cia2a is proposed to serve as a specific maturation factor for iron regulatory protein 1 (IRP1/ACO1), a cytosolic Fe-S protein regulating cellular iron homeostasis (**Figure 1A**).^18^ Notably, IRP1 does not end in a TCR motif and it is unknown what residues of IRP1 mediate its interaction with the Cia1-Cia2a complex.

Alignment of 21 Cia2a/b pairs with a subset of fungal (*Sc* and *Ct*) and plant (*Arabidopsis thaliana, At*) Cia2 sequences revealed that the TCR peptide binding Arg residue identified in *Ct* and *Sc* Cia2 is absolutely conserved in both Cia2a and Cia2b (**Figure 4A** and **S7**). We were surprised to see that the TCR binding residues are conserved in Cia2a proteins, since these proteins are proposed to be specific targeting factors for the TCR-less IRP1 protein.^18^ The conservation of TCR peptide binding residues in *Hs*Cia2a and its homologs suggests that Cia2a could facilitate maturation of clients terminating in a TCR motif.

**Figure 4.**
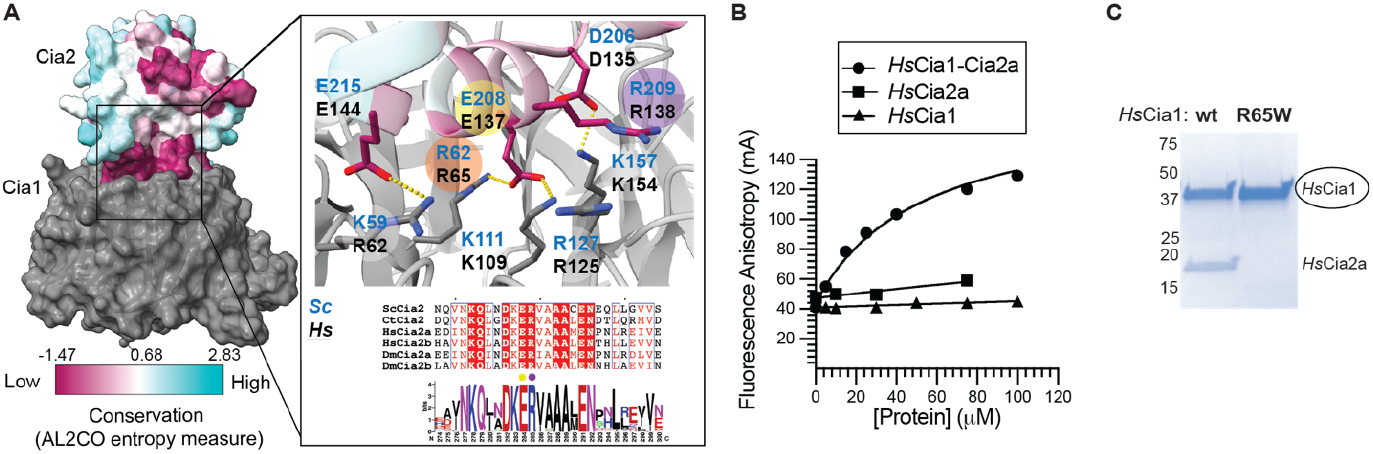
*Hs*Cia2a facilitates recognition of the TCR peptide. (**A**) The *Dm*Cia1-Cia2b complex (6TBN)^24^ is shown, with the Cia2b colored according to conservation in 47 Cia2 sequences, including two fungal (*Ct* and *Sc*) and three plant (*At*) Cia2s in addition to 21 Cia2a/b pairs. The inset shows a detailed view of the Cia1-Cia2 interface, a sequence alignment showing key Cia1-and TCR-binding region of Cia2, colored by conservation by ESPrint 3.0, and a WebLogo illustrating conservation of this region in 47 Cia2 sequences. See **Figure S7** for full alignment and details.^31, 32^ Orange and yellow circles highlight residues forming intersubunit salt bridge. Cia2’s TCR peptide binding Arg residue (purple circle) is adjacent to Cia1-binding glutamate (yellow circle). (**B**) FA assay monitoring interaction of FITC-4mer (0.1 µM) with the indicated protein(s) were completed as in **Figure 3A.** (**C**) Affinity copurification experiment comparing binding of *Hs*Cia2a prey to the WT *Hs*Cia1 or R65A variant (strep-tagged bait, circled).

To directly test this hypothesis, the *Hs*Cia1 and *Hs*Cia2a were purified, and their complex was isolated and utilized in the FA assay with the FITC-4mer probe (**Figure 4B**). We found that the *Hs*Cia1-Cia2a complex, but neither of the individual subunits, was able to bind the FITC-probe with an apparent *K*_*D*_ value of 48±11µM (**Table S2** and **Figure S8A-B**). Additionally, both the ^R125A^*Hs*Cia1-Cia2a and *Hs*Cia1-^R138A^Cia2a variants, analogous to the TCR peptide binding hotspot residues in the *Ct* homologs, disrupted binding to the FITC-probe (**Table S2** and **Figure S8A-B**). Altogether, these data indicated that the *Hs*Cia1-Cia2a complex can recognize clients terminating in TCR peptide.

The Cia2 sequence alignments also revealed the TCR peptide-binding residue (purple circle, **Figure 4A**) immediately follows a key residue mediating the Cia1-Cia2 interaction (yellow circle) via an intersubunit saltbridge.^24, 28^ The close proximity of TCR peptide-binding and Cia1-binding residues suggest a mechanism for coupling client recruitment to formation of the CTC. To investigate how defects in CTC formation could impact client recruitment, we focused on the ^R65W^*Hs*Cia1 variant recently identified as a pathogenic mutation implicated to a fatal neuromuscular disorder.^33, 34^ Based on the structure available for the *Dm* orthologs, R65 of *Hs*Cia1 contacts E137 of *Hs*Cia2a (**Figure 4A**), leading us to the hypothesis that the disease-causing R65W variant will disrupt complexation with *Hs*Cia2a and thus TCR peptide binding. Indeed, the ^R65W^Cia1’s interaction with *Hs*Cia2a was strongly disrupted (**Figure 4C** and **S8C**) without impacting Cia1’s thermostability (**Table S2**). Titration of the ^R65W^Cia1 variant into a solution containing *Hs*Cia2a and the FITC-4mer probe indicated this Cia1 variant results in a >10-fold lower affinity the TCR peptide with >10-fold weakened affinity (**Figure S8D**). Thus, this disease-causing variant affects CIA function by destabilizing the Cia1-Cia2 complex, thus decreasing the concentration of CTC available to maturate apo-clients.

## Discussion

Herein, we found that the TCR peptide binds at the interface of the Cia1-Cia2 interface. Computational and experimental assays identified two hotspot residues, corresponding to R127-*Sc*Cia1 and R209-*Sc*Cia2, each contributing ≥2 kcal/mol to binding energy. Our study clarifies the role of R127, which sits atop *Sc*Cia1’s third β-propeller domain. This residue was previously linked to CIA function, but its role was poorly defined.^27^ Our study also underscores the tight link between CTC formation and client recruitment, as R209 of *Sc*Cia2 lies within a conserved motif mediating Cia1-Cia2 interaction.^28^ Finally, the absolute conservation of TCR peptide binding residues in Cia2a proteins suggests an they play an unexpected role in recognizing clients ending in a TCR motif.

One key finding from our study is that Cia2 is directly involved in client identification. Early studies suggested that Cia2 interacts with clients, but subsequent reports argued the Cia2-client interaction is indirect, mediated by the other CTC subunits.^17-19, 24, 26^ These conflicting conclusions likely stem from reliance on qualitative coimmunoprecipitation assays to dissect these complex, multipartite interactions. Because perturbation of one CTC subunit is well-known to affect the others, qualitative cell-based assays can yield ambiguous or misleading data.^13, 15, 16, 18, 35^ Our data herein further underscores the pitfalls of relying exclusively on qualitative assays. The copurification data indicated the *Sc*Cia1 W201A and Y254A variants disrupt interaction with the TCR tripeptide without impacting Cia2 binding (**Figure 2** and **S3**). However, subsequent quantitative MST analysis showed these variants destabilize the Cia1-Cia2 complex, so their effect on TCR peptide binding must be indirect (**Table S1**). Our study underscores the need to rely on direct, quantitative *in vitro* assays when dissecting the complex interaction network driving CIA client recruitment.

Our study also clarifies Fe-S cluster trafficking pathway in the late stages of the CIA pathway. The fact that Nar1 competes with the Leu1 client for binding to the CTC suggests that Nar1 does not directly pass its cluster to clients. Instead, this data indicates Nar1 donates its cluster to the CTC then dissociates (**Figure 1C**).^21, 28^ Next, the Fe-S-bound CTC recruits and maturates an apo-client. This model predicts that Fe-S binding by Nar1 or the CTC will impact their interaction affinity and dynamics. Indeed, mutation of client Fe-S binding ligands can increase the stability of the client-CTC complex *in vivo*.^26, 36^ However, we and others have been unable to isolate the Nar1 Fe-S trafficking protein or the CTC in a fully a homogeneous, holo-state.^12, 21, 28, 37^ Future work will undoubtedly focus on how to prepare Fe-S cluster bound Nar1 or Cia1-Cia2 so that the quantitative assays developed herein can be applied to uncover how the trafficked Fe-S cluster affects these critical interactions.

Our discovery that binding of the TCR peptide requires the coordinated action of both Cia1 and Cia2 explains how the cell prevents individual subunits from competing with the CTC for binding apo-clients. The vital importance of Cia1-Cia2 complex formation for client recruitment is further emphasized by our investigation of the R65W *Hs*Cia1 variant implicated in a fatal, inherited neuromuscular disorder.^33, 34^ Although this variant did not affect the stability of *Hs*Cia1 *in vitro*, it did impact affinity for Cia2a and thus TCR peptide binding (**Figure 4C** and **S8C-D**). We suspect that the reported instability of the R65W variant *in vivo* likely results from its inability to be effectively incorporated into the CTC, leading to its posttranslational regulation as observed for other CTC subunits.^16, 38^

Finally, identifying peptide-binding hotspot residues unexpectedly revealed the metazoan Cia2 paralog, Cia2a, binds the TCR peptide. Undoubtedly, TCR binding residues of Cia2a allow it to recruit the Nar1 cluster trafficking protein via Nar1’s TCR motif. The proposal that Cia2a is a *bona fide* TCR peptide binding protein is supported by data from high throughput proteomics studies.^18^ Among the 15 proteins pulled down with Cia2a (those with ≥15 peptide matches), five - POLD1, NARFL, PPAT, RAF1, and MLF2 - end with a Trp or Phe residue. For comparison, 20% (n=44 proteins) of proteins immunoprecipitated with Cia2b terminate with these aromatic residues. These findings challenge the model that Cia2a is solely responsible for IRP1 maturation. Given Cia2a’s lower abundance in most tissues,^39^ we speculate Cia2a could also function as auxiliary general targeting factor, potentially facilitating CIA client maturation in cell types requiring high flux through the CIA system or under conditions of oxidative stress or iron limitation similar to the proposed roles of the bacterial ISC and SUF systems. Although further studies are needed to decipher the interplay between Cia2a and Cia2b within the cell, our work clearly shows that Cia2a likely plays a more general role in Fe-S protein maturation than previously hypothesized.

## Supporting information

Materials and Methods; Tables S1-S4 which include all measured dissociation constants, melting temperatures, and results from FTMap and PrankWeb analysis; Figures S1-S8 which include uncropped SDS-PAGE gels and western blots of affinity copurification experiments included in the main figures, including input samples and controls, multiple sequence alignments, and representative FA and MST data.

## Acknowledgement

This work was supported by the NSF-GRFP (M.M.), the NIH R01GM 121673 (to D.L.P), and by the Boston University Undergraduate Research Opportunities Program (C.C.Q).

