## supporting information for "The Cia1 and Cia2 subunits of the CTC mediate recognition of apo-FeS proteins with a C-terminal targeting complex recognition motif"

##### Affiliations: Department of Chemistry, Boston University, Boston, MA USA.

##### ^‡^these authors contributed equally to this work

Includes:

Materials and Methods

Supporting Tables S1-S4

Supporting Figures S1-S8

**Materials and Methods**

##### *Plasmid construction*

For *C. thermophilum* (*Ct*) proteins, the *E. coli* codon optimized gene corresponding to *Ct*Cia1 (uniprot: G0RYD4_CHATD) and *Ct*Cia2 (uniprot: G0SBV3_CHATD) were synthesized (GenScript) and cloned into pETDuet-1 (Millipore Sigma). *CIA1* was cloned into multiple cloning site one with N-terminal His_6_-tag and human rhinovirus 3C protease cleavage site before the Cia1 coding sequence (^His^*Ct*Cia1)*.* *Ct*Cia2 was cloned into multiple cloning site two such that the expressed protein has an N-terminal Strep-tag and tobacco etch virus (TEV) protease cleavage site (^Strep^*Ct*Cia2).

For *H. sapiens (Hs)* proteins, the *E. coli* codon optimized gene for *Hs*Cia1 (uniprot: CIAO1_HUMAN) was cloned into pETDuet-1 with a sequence encoding an N-terminal His_8_-TEV-Strep tag (^DT^*Hs*Cia1, for double-tagged *Hs*Cia1). For *Hs*Cia2a (uniprot: CIA2A_HUMAN), the codon optimized gene for amino acids 28-160 was cloned into pGEX-4T (Millipore Sigma) creating a truncated *Hs*Cia2a with an N-terminal His_8_-GST tag (^GST^*Hs*Cia2a).

All site-directed mutants were introduced via Q5 mutagenesis (New England Biolabs) according to manufacturer’s instructions and verified via DNA sequencing.

##### *Protein purification*

For *S. cerevisiae* (*Sc*) proteins, Cia1 [uniprot: CIAO1_YEAST; with N-terminal His_6_-tag (^His^Cia1) or N-terminal His_6_-TEV-Strep double tag (^DT^Cia1)], Cia2 [uniprot: YHS2_YEAST; with N-terminal Strep-tag and C-terminal His_6_-tag (double tagged,^DT^Cia2), the ^Δ102^Cia2 (in which the N-terminal 102 amino acids are deleted and there is a C-terminal His_6_-tag], ^SUMO^Met18 [uniprot: MET18_YEAST; with N-terminal His_6_-SUMO tag], ^SUMO^Nar1 [uniprot: NAR1_YEAST; with N-terminal His_6_-SUMO tag], ^His^Leu1 [uniprot: LEUC_YEAST; with N-terminal His_6_-TEV tag] and SUMO peptide carrier (SPC, His_6_-SUMO) proteins ^SPC^Leu1 (last 21 amino acids of *Sc*Leu1 attached to the C-terminus SPC) and ^SPC^Nar1 (last 10 amino acids of *Sc*Nar1 attached to the C-terminus of SPC) were expressed and purified as previously described.^1-3^

To purify the *Sc*Cia1-Cia2 complex, ^DT^Cia2 (~4 mg) was mixed with 10 mg of ^His^Cia1. After incubation for 1h at 4°C, the mixture was batched absorbed to Streptactin XT Superflow (IBA) resin (2 mL) equilibrated with Tris buffer [50 mM Tris-HCl (pH 8.0), 100 mM NaCl, 5 % glycerol, 5 mM BME]. After 1h, the resin was collected, washed with 15-20 CV of Tris buffer to remove any Cia1 not involved in formation of the Cia1-Cia2 complex, and eluted with Tris buffer supplemented with 30 mM D-Biotin (IBA) and 40 mM NaOH (to adjust pH to 8.0). Protein containing fractions were combined and exchanged into Tris buffer via a PD10 gel filtration column (Fisher Scientific). The protein was then concentrated to 2-3 mg/mL using 10 kD Amicon ultra centrifugal filter unit (MilliporeSigma) and stored at –80°C.

For copurification of the *Ct*Cia1-Cia2 complex, the co-expression plasmid was transformed in BL21(DE3) (New England Biolabs) grown in LB with 100 µg/mL carbenicillin (LB/Carb) at 37°C to an OD_600_ of 0.6−0.8. Isopropyl β-D-1-thiogalactopyranoside (IPTG, 1 mM) was added and the cells were collected 3.5-4 h later.

For purification at 4 °C, cell paste (10-20 g) was resuspended in 100-200 mL of NaP_i_ buffer [50 mM Na_2_HPO_4_ (pH 8.0), 100 mM NaCl, 5% glycerol, and 1 mM DTT] supplemented with 1 mM PMSF (GoldBio), protease inhibitor tablet (Thermo Scientific), 4 kU DNase nuclease (Thermo Scientific), and rLysozyme (45 kU/g of cell, Millipore Sigma). Cells were disrupted by microfluidization. The crude extract was centrifuged, and the soluble fraction was added to 4-8 mL of Streptactin XT Superflow resin. The column was washed with 15-20 column volumes (CV) of NaP_i_ buffer and eluted with NaP_i_ buffer supplemented with 30 mM D-Biotin and 40 mM NaOH (pH 8.0). Protein containing fractions were combined and concentrated using 10 kD Amicon ultra centrifugal filter units (Millipore Sigma) and dialyzed overnight against NaP_i_ buffer. The protein (5-10 mg/mL) was stored at –80°C. The *Ct*Cia1-Cia2 variants were purified as described for the wild-type complex except the cells (~7 g) were lysed by the addition of CelLytic Express (5.0 g, Sigma Aldrich) and exchanged into NaP_i_ buffer via a PD-10 gel filtration column equilibrated in NaP_i_ buffer.

Plasmids encoding ^DT^*Hs*Cia1 and ^GST^*Hs*Cia2a, WT or variants, were separately transformed into BL21(DE3) and grown as described for *Ct*Cia1-Cia2 complex, except 0.5 mM IPTG was added. For purification of the *Hs*Cia1-Cia2a complex, the cell pastes from *Hs*Cia1 and *Hs*Cia2a overexpression were mixed in a 1:2 ratio, with totaling ~15 g of cell paste. Hepes buffer [50 mM Hepes (pH 8.1), 100 mM NaCl, 5% glycerol, 1 mM DTT] was added (75 mL) along with a protease inhibitor cocktail tablet (Thermo Scientific), 1 mM PMSF (Goldbio), 10 kU DNase nuclease (Thermo Scientific), and 0.5% octylthioglucoside (GoldBio). Cells were disrupted via sonication. After centrifugation, the soluble extract was added to Streptactin XT Superflow resin (2 mL, IBA). The column was washed with 15 CV of Hepes buffer, 15 CV of Hepes buffer with 150 mM NaCl, and eluted with Hepes buffer supplemented with 40 mM D-Biotin (IBA) and 50 mM NaOH (pH 8.1). Protein containing fractions were combined, concentrated using 30 kD Amicon ultra centrifugal filter units (MilliporeSigma) and exchanged into Hepes buffer via a PD-10 gel filtration column.

For the separate purification of *Hs*Cia1 and *Hs*Cia2a, the ^DT^*Hs*Cia1 was purified as described for the *Hs*Cia1-Cia2a complex except cells expressing Cia2a were not included during cell resuspension. For purification of ^GST^*Hs*Cia2a, 8-10 g of cell paste was resuspended with 40-50 mL of Hepes buffer with a protease inhibitor cocktail tablet (Thermo Scientific), 1 mM PMSF (Goldbio), and 10 kU of DNase nuclease (Fisher Scientific), and 0.5% octylthioglucoside (GoldBio). The cells were disrupted via sonication. After centrifugation, the soluble extract was added to Ni-NTA resin (2 mL, GoldBio). The column was washed with 15 CV of Hepes buffer with 5 mM imidazole, 15 CV of Hepes buffer with 10 mM imidazole, and eluted with Hepes buffer with 250 mM imidazole. Protein containing fractions were combined, concentrated using 30 kD Amicon ultra centrifugal filter units (MilliporeSigma) and exchanged into Hepes buffer via a PD-10 gel filtration column.

To remove the GST-tag from *Hs*Cia2a, TEV-protease cleavage was performed by mixing 10 mg of protein with 1 mg of TEV-protease in the presence of 5 mM DTT. The mixture was gently rocked overnight at 4°C then applied to a 2 mL Streptactin XT Superflow column and purified as described for *Ct*Cia1-Cia2 except the washing was reduced to 5 mL of Hepes buffer supplemented with 150 mM NaCl.

*Affinity copurification assay*

Assays were carried out as described.^1-3^ Briefly, a ~200 µg of a strep tagged bait protein was mixed with equimolar amount the each indicated prey protein in ~900 µL. After a 1h incubation at 4°C, the mixture was batch absorbed to 200 µL of streptactin XT Superflow resin (IBA). After 1h, resin was collected, washed, and eluted with biotin containing buffer. Input and elution fractions were analyzed by SDS-PAGE and western blot (His-Tag (27E8) Mouse mAb HRP Conjugate, Cell Signaling Technologies). When ^DT^*Sc*Cia2 was the bait, then ^His^*Sc*Cia1, ^SUMO^*Sc*Met18, ^SUMO^*Sc*Nar1, ^His^*Sc*Leu1, ^SPC^Leu1 and/or ^SPC^Nar1were used as the prey proteins, as indicated. When ^DT^*Sc*Cia1 was the bait, ^SUMO^Nar1 was the prey. When *Ct* proteins were analyzed, *Sc*Leu1 or *Sc*Met18 were the prey, and the copurified complex of ^His^*Ct*Cia1- ^Strep^*Ct*Cia2 was utilized as the strep-tagged bait. To test for interaction of *Hs* proteins, ^DT^*Hs*Cia1 was the strep-tagged bait and *Hs*Cia2 (GST cleaved) was the prey. In all figures, the strep-tagged bait protein is indicated by a circle.

Several controls were routinely analyzed in parallel with each copurification experiment. First, SDS-PAGE analysis of input samples was used to ensure any variants being compared are present at similar concentrations, as shown in supporting information figures. When a variant was expected to disrupt an interaction, a positive control using the wild-type proteins was included. A negative “no bait” control monitored for any nonspecific interaction of the prey proteins with the resin. In assays utilizing ^SPC^Leu1 or ^SPC^Nar1 as prey proteins, additional two negative controls included: i) a sample omitting any SPC-prey to ensure that any low molecular weight proteins detected in the SDS-PAGE gel or western blot are associated with the SPC-prey and not from proteolysis of other assay components because many also contain a His-tag; and ii) a control with the His_6_-SUMO peptide carrier (SPC) without addition of any C-terminal TCR peptide sequences to monitor for any nonspecific interaction of the SPC tag with the resin or the CTC components. All these controls are included in uncropped western and SDS-PAGE gels included in the supporting information figures.

### *Computational prediction of TCR-peptide binding site*

The *Dm*Cia1-Cia2B (PDB: 6TBN)^4^ was analyzed using the publicly available webservers (ftmap.bu.edu) and PrankWeb (prankweb.cz).^5, 6^ For FTMap analysis, the inside of Cia1’s central cavity was masked to direct probes to viable protein-protein interaction sites on the protein surface rather than inside the beta propeller cavity. The PyMOL Molecular Graphics System was used to visualize the results and create the figures.

*Fluorescence anisotropy assays*

The 4-mer peptide (HQDW; GenScript) or 21-mer peptide (FDNVPKRKAVTTTFDKVHQDW; ABM) with N-terminal FITC attached to a 1,6 aminohexanoic acid spacer was dissolved in 0.125% NH_4_OH. The concentration of the FITC-HQDW peptide was determined from its FITC extinction coefficient (493 nm, 70,000 M^-1^ cm^-1^).

Fluorescence anisotropy assays were performed in 96-well polypropylene black plates (Corning Costar, #3356). FITC-peptide probe (0.1 μM) diluted into buffer supplemented with 0.1 mg/mL BSA and the indicated CTC subunit(s) (0-100 µM) were mixed in a final volume of 200 µL and incubated at 25 °C for 1h. For *Sc* proteins, a Tris-FA buffer [50 mM Tris-HCl (pH 8.0), 100 mM NaCl, 5% glycerol, 5mM BME] was used. For *Ct* proteins, a NaP_i_-FA buffer [50 mM Na_2_HPO_4_ (pH 8.0), 100 mM NaCl, 5% glycerol, 5 mM BME] was used. For the *Hs* proteins, a Hepes-FA buffer [50 mM Hepes (pH 8.1), 100 mM NaCl, 5% glycerol, 5mM BME] was used and the GST tag was removed from *Hs*Cia2a. All other *Sc* and *Ct* proteins were analyzed without cleaving any of their purification tags.

The fluorescence anisotropy was determined using a SpectraMax5 plate reader (Molecular Devices) at 25°C with excitation at 488 nm and emission at 520 nm employing a 515 nm cutoff filter in high-sensitivity mode with 100 reads/well (on-calibration, normal speed, and column priority). The fluorescence anisotropy was calculated using **Equation 1:**

$$A =\left( \frac{I_{para}-I_{perp}}{I_{para}+2\cdot I_{perp}} \right)$$

where *A* is the anisotropy, *I_par_*_a_ and *I_perp_* are the intensity of the emitted light in the parallel plane and perpendicular plane, respectively. The background anisotropy values were measured before adding the FITC-peptide and subtracted from the anisotropy determined after the addition of the probe. The total fluorescence of each sample was also measured using the same excitation and emission wavelength in auto-sensitive mode with 6 flashes per read (read from top, on-calibration column priority).

To determine the molar concentration of the Cia1-Cia2 complex, a 1:1 complex between Cia1 and Cia2 was assumed to convert the concentration determined by the Bradford assay using BSA as the standard. To ensure that the protein stock concentrations for WT and tested variants are the same, A_280_ was compared. Additionally, an SDS-PAGE analysis of samples was performed to ensure that impurities and stoichiometry for Cia1 and Cia2 in complexes were similar in all samples.

The background subtracted anisotropy values (*A*) were plotted versus protein concentration and data was fitted to a hyperbolic binding model (**Equation 2**) using GraphPad Prism 7.

$$A = \left( A_{max}-A_{min} \right)\cdot\left( \frac{x}{x+K_{D}} \right)+A_{min}$$

where *A_max_* is the maximum anisotropy, *A_min_* is the minimum anisotropy, *x* is the concentration of the Cia1-Cia2 complex, and *K_D_* is the dissociation constant. For all variants, the *A_max_* value was constrained to the value observed with wild-type protein (180 mA).

For comparison of FITC-4mer and FITC-21mer, the background subtracted anisotropy data were normalized using **Equation 3**

$A_{norm}=\frac{A_{raw}-A_{f.min}}{A_{f.max}-A_{f.min}}$

where *A_raw_* is the background subtracted anisotropy, *A_f.min_* and *A_f.max_* are the minimum and the maximum fitted anisotropy values determined by fitting the raw data to Equation 2. The normalized anisotropy was plotted against the protein concertation and fitted to Equation 2, contraining the *A_max_* to 1, and *A_min_* was to 0.

The change in binding energy (ΔΔG, in kcal/mol) obtained from the alanine scanning mutagenesis was calculated using **Equation 4.**

$\Delta\Delta G = -RT\ln\left( \frac{K_{D}^{WT}}{K_{D}^{mut}} \right)$

where *K_D_^WT^* and *K_D_^mut^* are the dissociation constants for wild-type protein and its mutant, respectively, T= 298K, R = 0.001987 kcal•mol^-1^•K^-1^.

To determine the effect of R65W mutation on *Hs*Cia1-Cia2 complex formation, FITC-peptide probe (0.2 μM), BSA (0.2 mg/mL), and *Hs*Cia2A (20 µM, GST tag removed) were diluted in HEPES-FA buffer in a final volume of 100 µL. After incubation at 25 °C for 1h, a 100 µL aliquot of *Hs*Cia1 or ^R65W^Cia1 (both with N-terminal His_8_-TEV-Strep tag) was added such that the final Cia1 concentration ranged from 0-40 µM in a final volume of 200 µL. The mixture was incubated at 25°C for 1h. The FA data were collected and analyzed as described for the *Sc* and *Ct* samples except when data was fit to Equation 2, the *A_max_* was constrained to value observed with wt *Hs*Cia1 (90).

##### *Melting temperature determination*

Nano Differential Scanning Fluorimetry (NanoDSF) analysis was performed using a Prometheus NT.48 instrument (NanoTemper). *Ct*Cia1-Cia2 was diluted to 10 µM in NaP_i_-FA buffer and *Sc*Cia1-Cia2 was diluted to 10 µM in Tris-FA buffer. The diluted samples were centrifuged and introduced to capillaries. Capillaries were heated from 20-95°C with a 1°C per min ramp rate. The fluorescence intensity was measured with an excitation wavelength of 280 nm, excitation power (28% for *Ct*, 35% for *Sc,* and 20% for *Hs)*, and emission wavelengths of 330 and 350 nm. The melting temperature (T_m_) of proteins was determined from the first derivative of the ratio of the 350 and 330 nm fluorescence intensities.

For *Hs*Cia1 proteins, Realplex Mastercycler RT PCR instrument (Eppendorf) was used. The assay mixture (20 µL) consisted of 32 µM *Hs* ^DT^Cia1 (double tagged with His_8_-TEV-Strep tag) in Hepes-FA buffer and SYPRO orange dye (5X, ThermoFisher Scientific) were added to a 96-well PCR plate and fluorescence at 520 nm was monitored as the temperature was increased from 20-95°C with a 1°C per min ramp rate. The T_m_ values were calculated as the peak of the first derivative of the melting curve (fluorescence at 520 nm vs temperature).

*Bioinformatic analysis*

To evaluate the frequency of Trp or Phe residue within the TCR motif at the C-terminus of CIA client proteins the catalog of CIA client protein from eukaryotic organisms in Marquez *et al.*^1^ was utilized and further extended by adding homologs for *G. gallus*, *C. thermophilum*, *C. albican*, *G. max*, *O. sativa*, *A. carolinensis*, *D. discoideum*, *C. neoformans*, *C. reinhardtii*, *S. lycopercicum*, and *H. vulgaris.* In total 23 organisms were used. The sequences were identified by NCBI BlastP searches using yeast and human sequences as queries. The sequences where all residues in C-terminal six positions were the same were removed and then the frequency of amino acids at the C-terminal position was calculated.

The surface of the CTC was colored by conservation in ChimeraX based on Clustal Omega alignment of Cia1, Cia2b, and Met18 homologs from *H. sapiens, S. cerevisiae, S. pombe, D. rerio, X. laevis, D. melanogaster, A. thaliana,* and *C. elegans.*

The Cia2a/b sequences were identified by NCBI BlastP searches using yeast and human sequences as queries. The analysis included the following organisms: *Homo sapiens (Hs), Drosophila melanogaster (Dm), Danio rerio (Dr), Anolis carolinensis (Ac), Xenopus laevis (Xl), Cavia porcellus (Cp), Loxodonta africana (La), Ornithorhynchus anatinus (Oa), Xiphophorus maculatus (Xm), Anopheles gambiae (Ag), Aedes aegypti (Aa), Apis mellifera (Am), Latimeria chalumnae (Lc), Mustela putorius furo (Mp), Tetrahymena thermophila (strain SB210) (Tt), Ciona intestinalis (Ci), Hydra vulgaris (Hv), Branchiostoma floridae (Bf), Nematostella vectensis (Nv), Meleagris gallopavo (Mg),* and *Aquila chrysaetos chrysaetos (Acc).* Cia2 sequences from *Saccharomyces cerevisiae (Sc), Chaetomium thermophilum (strain DSM 1495 / CBS 144.50 / IMI 039719) (Ct),* and *Arabidopsis thaliana (At)* were also included in the alignment*.* Multiple sequence alignment was performed with Clustal Omega, followed by the identification of conserved regions in ESPript 3.0 and WebLogo.^7, 8^

##### *Microscale thermophoresis (MST) assay*

Either ^His^*Sc*Cia1 or ^Δ102^*Sc*Cia2 were labeled using the RED-NHS (Cia1) or RED-MALEIMIDE (Cia2) Monolith NT Protein Labeling Kit according to manufacturer instructions (NanoTemper Technologies) to create the labeled “target” protein. The unlabeled *Sc* binding partner (0.15 nM-8.1 µM) was incubated with 20 nM labeled target protein in 50 mM Tris-HCl (pH 8.0), 100 mM NaCl, 5% glycerol, 5 mM BME, 50 mM Arg, and 0.05% Tween-20. Samples were loaded onto standard or premium capillary tubes (NanoTemper Technologies). After 5 min incubation at room temperature, the assay was performed using a NanoTemper Monolith NT.115 instrument at 20-60% excitation power and medium MST power. Analysis of the MST data was completed using the manufacturer’s software package. Briefly, the ligand dependent change in the microscale thermophoresis (*F_norm_*) upon IR irradiation was first normalized (Δ*F_norm_*) by subtracting the *F_norm_* value observed in the absence of ligand (the unbound state) from the *F_norm_* value at each concentration of ligand. The normalized MST response (Δ*F_norm_*) was then fit to a quadratic binding model to determine the *K_D_* value. For presentation of data in figures where multiple datasets are presented on the same graph, the Δ*F_norm_* was normalized by the amplitude of the binding curve to present data as fraction bound.

**Table S1. *K_D_* values for the *Sc*Cia1-Cia2 interaction determined by MST**

| Unlabeled component | Labeled component | *K_D_*, nM^1^ |
| --- | --- | --- |
| WT Cia2 | WT Cia1 | 24±8.4 (n=3) |
| ^E208A^Cia2 | WT Cia1 | 1220 ±0.4 (n=2) |
| ^R209A^Cia2 | WT Cia1 | 72 ±15 (n=3) |
| WT Cia1 | ^Δ102^Cia2 | 40 ±20 (n=3) |
| ^W18A^Cia1 | ^Δ102^Cia2 | 1620 ±110 (n=3) |
| ^R62A^Cia1 | ^Δ102^Cia2 | 1210 ±210 (n=3) |
| ^E104A^Cia1 | ^Δ102^Cia2 | 1.7±0.3 (n=2) |
| ^E109A^Cia1 | ^Δ102^Cia2 | 77 ±66 (n=2) |
| ^K111A^Cia1 | ^Δ102^Cia2 | 6.4±2.6 (n=3) |
| ^R127A^Cia1 | ^Δ102^Cia2 | 12±0.8 (n=3) |
| ^K129A^Cia1 | ^Δ102^Cia2 | 378 ±52 (n=2) |
| ^W132A^Cia1 | ^Δ102^Cia2 | 1180 ±220 (n=2) |
| ^V148A^Cia1 | ^Δ102^Cia2 | 0.9±0.5 (n=3) |
| ^D155A^Cia1 | ^Δ102^Cia2 | 133 ±17.5 (n=2) |
| ^W201A^Cia1 | ^Δ102^Cia2 | 1000 ±660 (n=2) |
| ^Y254A^Cia1 | ^Δ102^Cia2 | 315 ±29 (n=3) |
| ^E297A^Cia1 | ^Δ102^Cia2 | 38000 ±3300 (n=3) |
| ^N299A^Cia1 | ^Δ102^Cia2 | 4100 ±460 (n=3) |

^1^ The number of replicates (n) is indicated. The error is the standard deviation of the independent measurements.

**Table S2. Apparent dissociation constants for TCR peptide binding Cia1 and Cia2 variants.**

| Organism | Cia1-Cia2 variant | *K_D_*, μM^1^ | ΔΔG^2^, kcal/mol | T_m_,^3^ °C | ∆T_m_,^2^ °C |
| --- | --- | --- | --- | --- | --- |
| *Sc* | WT Cia1-Cia2 | 86±0.4 (n=2) | -- | 53.3 | -- |
|  | ^R127A^Cia1-Cia2 | >10000 | >3.1 | 57.8 | +4.5 |
| *Ct* | WT Cia1-Cia2 | 25±3.2 (n=3) | -- | 69.5 | -- |
|  | ^R163A^Cia1-Cia2 | 1000±210 (n=2) | 2.2 | 75.2 | +5.7 |
|  | ^E145A^Cia1-Cia2 | 49 | 0.4 | 67.2 | -2.3 |
|  | ^K165A^Cia1-Cia2 | 52±0.0 (n=2) | 0.4 | 69.6 | +0.1 |
|  | ^K147A^Cia1-Cia2 | 440 | 1.7 | 69.1 | -0.4 |
|  | Cia1- ^R177A^Cia2 | 780±62 (n=2) | 2.0 | 69.5 | 0.0 |
|  | Cia1- ^Q171A^Cia2 | 230±44 (n=2) | 1.3 | 69.5 | 0.0 |
| *Hs* | WT Cia1-Cia2A | 48±11 (n=4) | -- | ND^4^ | ND |
|  | Cia1- ^R138A^Cia2A | 270±7.0 (n=2) | 1.0 | ND | ND |
|  | ^R125A^Cia1- ^R138A^Cia2A | 670±310 (n=2) | 1.6 | ND | ND |
|  | ^R125A^Cia1-Cia2A | 1060±720 (n=2) | 1.8 | ND | ND |
|  | WT Cia1 | 17±0.86 (n=2)^5^ | -- | 47.9^6^ | -- |
|  | ^R125A^Cia1 | ND | ND | 50.6^6^ | +2.7 |
|  | ^R65W^Cia1 | 210±87 (n=2)^5^ | 1.5 | 47.9^6^ | 0.0 |

^1^ The apparent dissociation constant for the FITC-4mer probe was measured via the FA assay. The number of replicates (n) is indicated along with the standard deviation of the two or more independent measurements.

^2^ Relative to WT protein from the same organism.

^3^ T_m_ values listed correspond to melting temperatures determined by nanoDSF unless otherwise noted

^4^ ND, not determined

^5^Since the ^R65W^Cia1-Cia2a complex could not be isolated, the apparent *K_D_* was determined by titrating *Hs*Cia1 (WT or R65W variant) into a solution containing 0.1 µM FITC-4mer probe and 10 µM *Hs*Cia2a.

^6^T_m_ values was determined using Thermofluor assay.

**Table S3. Consensus Sites (CSs) identified by FTMap**

| **CS** | **Probe**  **Cluster^1^** | **Probes (total #)^2^** | ***Dm*Cia1^3^ residues** | ***Dm*Cia2b residues^3^** |
| --- | --- | --- | --- | --- |
| 1 | CC1 | BDY, CHX, DFO, DME, THS, ADY, AMN, BEN, BUT, ETH, PHN, ACD, EOL, URE, ACN, ACD, ACT (15) | D273, R253 | K109, V110, T111, V112, R98, A130, K132 |
| **2** | **CC2** | CAN, ADY, AMN, BEN, ETH, ACD, ACT, BDY, CHX, DFO, DME, PHN, THS, BUT (14) | - | L94, T90, L129, Q128, D131, R134, V135, A138 |
| 1 | CC3 | ACD, URE, CAN, EOL, ADY, AMN, BDY, BEN, DFO, ETH, BUT, PHN, ACT (13) | D274, Q304, D305, D324, R16 | R107, F108, P106, K109, V110, K132 |
| 1 | CC4 | BUT, ADY, DME, EOL, CAN, AMN, DFO, THS (8) | S252, R253, T216, A254, D214 | - |
| 1 | CC5 | ACD, AMN, URE, URE, BEN, ETH (6) | R16 | V110, L101, R97, V98, K132, V135, L139, A136 |
| 1 | CC6 | BEN, CHX, EOL, ADY, PHN(5) | Q304, D324, R16, G15 | P106, L101 |
| - | CC7 | BUT, EOL, PHN, DFO, URE(5) | D214, S195, T196 | L123, K127, A130 |
| **2** | **CC8** | DME, ACT, BDY, BEN, PHN(5) | R123, E105, K107 | R134, A138, A137, N141 |
| - | CC9 | ACT, CAN, DFO, EOL(4) | - | N69, I68, R67, S74, V75, H76 |
| 1 | CC10 | THS, DME, ACD(3) | Q304, D305, D324 | F108, K109, V110, R97 |
| - | CC11 | BUT, BDY, DME(3) | Q304, D325, D324, R16, G15, E34 | - |
| - | CC12 | ETH, ACN(2) | R16, E34 | A136, L139 |
| 2 | CC13 | AMN(1) | - | R97, L129, A130, D131, K132 |

**^1^** A probe cluster is a group of minimized probe conformations, grouped based on their structural similarities and energy levels. A consensus site (CS) is a 'hot spot' identified by overlapping these probe clusters within 4 Å, representing regions with a high probability of binding interactions.

^2^ The identities of individual probes found at each CCs are listed with 3 letter abbreviations and the total number of probes found at each CC is listed in parentheses. The aromatic probes are underlined. The abbreviations are as follows: ETH – ethane, EOL – ethanol, THS – Isopropanol, BUT – tert-Butanol, ACN – acetonitrile, AMN – methenamine, DFO – N, D-dimethylformamide, DME – dimethyl ether, BDY – benzaldehyde, BEN – benzene, CHX – cyclohexane, PHN – phenol, ACD – acetamide, ACT – acetone, AFY – acetaldehyde, URE – urea.

^3^ Arginine residues of *Dm*Cia1 and *Dm*Cia2 that correspond to the TCR peptide-binding hotspot residues identified for the *Ct* proteins are highlighted in yellow.

**Table S4. Sites identified by PrankWEB analysis**

| **Site** | **Score^1^** | **Conservation^2^** | ***Dm*Cia1 residues^3^** | ***Dm*Cia2b residues^3^** |
| --- | --- | --- | --- | --- |
| **1** | 6.93 | 1.639 | I16, Y256, Q304, D305, D324, E34 | L101, P106, P108, K109, V110, A130, K132, V135, A136, L139, R97, W98 |
| **2** | 6.7 | 3.215 | E105, R123 | Q128, L129, R134, V135, A137, A138, N141, L144, T90, L91, L94 |
| 3 | 3.18 | N/A | H157, K160, I162, A176, T187, H225, Q238, T239, W241 | -- |
| 4 | 2.95 | 0.083 | V155, W156, P158, I200, D201, F202, V258, S259, W260, K262, Q310, W311, P313, R66 | -- |
| 5 | 1.9 | 3.39 | -- | H119, S121, V125, Q128, H85, C86, S87, T90 |
| 6 | 1.88 | 1.325 | S110, W111, A21, W22, P24, Q310, W311, P313, R66, W67, P69 | -- |
| 7 | 1.72 | 1.263 | H23, V28, I40, K51, G71, K88, F93 | -- |
| 8 | 1.14 | 3.599 | -- | E50, H51, T82, I83, S87 |
| 9 | 1.12 | 1.414 | W111, S114, G115, V132, G134, F138, P69, C70, Y73 | -- |
| 10 | 0.71 | 1.987 | E100, H102, W128, W130, T83, L99 | -- |

^1^ Sites ranked according to the score assigned by the PrankWEB,^6^ incorporates the properties of protein residues including hydrophobicity, charge, and size. Overall, the analysis generates a score indicating the likelihood of a given pocket on the protein surface being a ligand-binding site.

^2^Conservation of residues lining the identified cavities as scored by PrankWEB. A higher value corresponds to a higher degree of conservation.

^3^ Arginine residues of *Dm*Cia1 and *Dm*Cia2 that correspond to the TCR peptide-binding hotspot residues identified for the *Ct* proteins are highlighted in yellow.

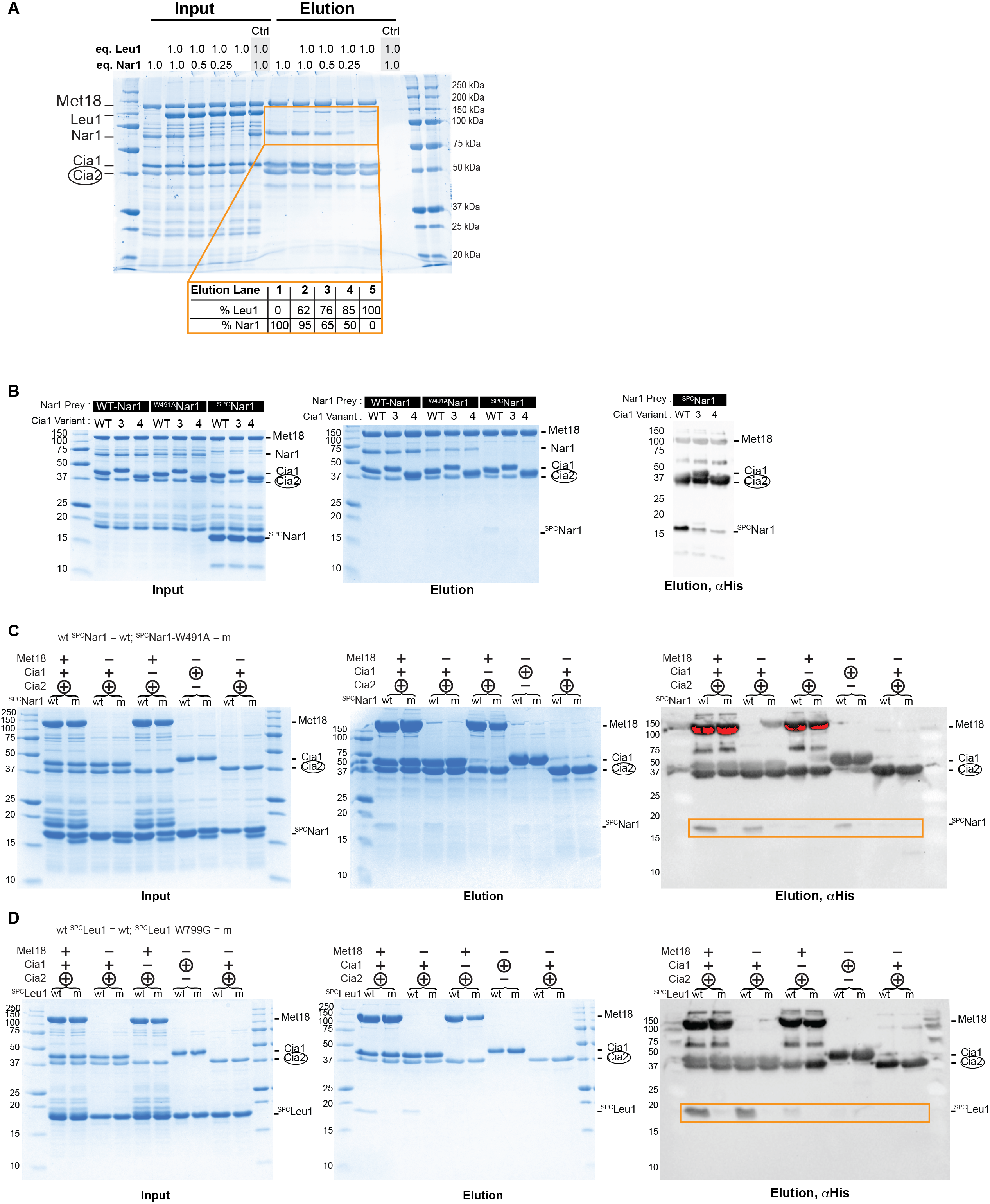

**Figure S1**. *Sc*Nar1’s interaction with the *Sc*CTC. (**A**) Uncropped input and elution gels related to **Figure 1A**. ^DT^Cia2 bait (circled) was mixed with ^SUMO^Met18, ^His^Cia1, ^His^Leu1, and increasing amounts of ^SUMO^Nar1, ranging from 0.25 to 1.0 molar equivalents relative to Leu1. After purification of the resulting complexes using streptactin resin, the input and elution samples, along with a negative control (ctrl) in which the ^DT^Cia2 bait was omitted, were analyzed by SDS-PAGE gel. The intensity of Leu1 and Nar1 bands in the elution fractions (yellow box) were determined by densitometry and ratioed against lane without any competitor, elution lane 1 for Nar1 or elution 5 for Leu1. (**B**) The indicated *Sc*Nar1 construct (WT Nar1, ^W491A^Nar1, or ^SPC^Nar1) was mixed with the indicated ^His^Cia1 variant (numbered as in **Figure 2D**) in the presence of ^SUMO^Met18 and strep-tagged Cia2 (bait protein, circled). The mixture was passed through a streptactin column and input and elution fractions analyzed via SDS-PAGE or western blot, for the His-tag on the SPC. Note that the CTC subunits all also contain a His tag, but they are differentiated from ^SPC^Nar1 based on its distinctive ~20 kDa molecular weight and absence of corresponding band when ^SPC^Nar1 is omitted. (**C**) and (**D)** Uncropped input and elution gels related to **Figure 2C**. The strep-tagged bait protein (circled) was mixed with the indicated CTC subunits and ^SPC^Nar1 (**C**) or ^SPC^Leu1 (**D**). The input and elution fractions from streptactin purification were analyzed by SDS-PAGE and western blot. Proteins were identified by comparison with migration of each purified recombinant protein. Region of western blot shown in **Figure 2C** are boxed.

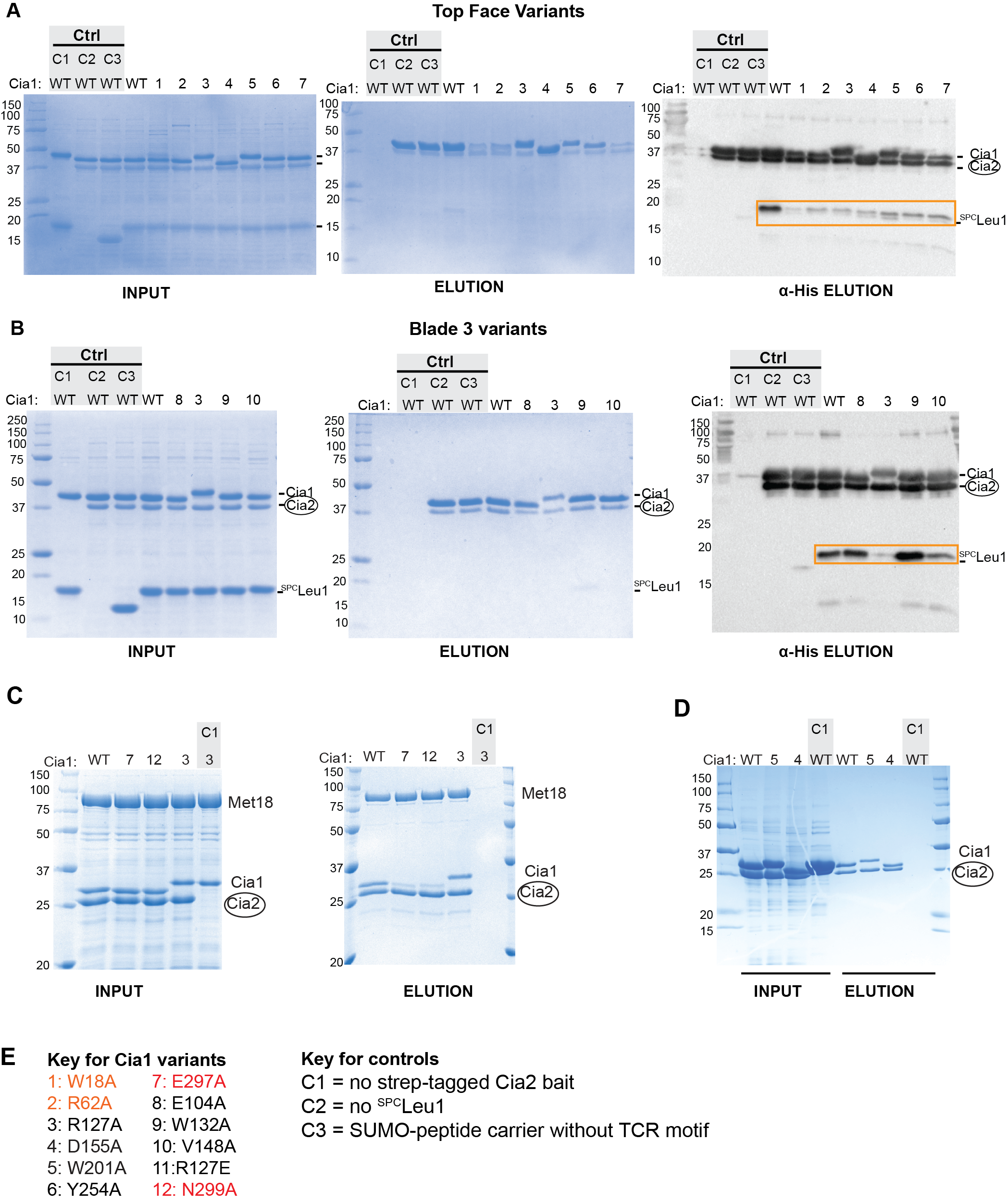

**Figure S2.** Top face of *Sc*Cia1 is important for binding *Sc*Cia2 and the TCR motif. (**A**) and (**B**) Uncropped SDS-PAGE gels (input and elution) and western blots (elution only). Orange box corresponds to data shown in **Figure 2E.** The ^His^Cia1 variants are numbered according to the key in this figure’s panel **E** and as in **Figure 2D**. Three control (ctrl) lanes are also shown. In C1, the strep-tagged Cia2 bait (^DT^Cia2) was omitted to monitor for any nonspecific interaction of bait proteins (^SPC^Leu1 and/or ^His^Cia1) with the streptactin resin. In the C2 samples, ^SPC^Leu1 was excluded to ensure any low molecular weight species observed in the SDS-PAGE or western blot are associated with ^SPC^Leu1. Although both ^His^Cia1 and ^DT^Cia2 have a His-tags, their higher molecular weight allows them to be distinguished from ^SPC^Leu1. In the C3 control samples, the SUMO-peptide carrier (SPC only, without the C-terminal tail of Leu1) was included to monitor for nonspecific interactions with the SUMO tag. (**C**) Affinity copurification of ^His^Cia1 (WT or variant, as indicated) and ^SUMO^Met18 with the ^DT^Cia2 bait (circled). The gel also includes negative control (C1) in which the ^DT^Cia2 bait was omitted. (**D**) Affinity copurification experiments as **S2C**, except ^SUMO^Met18 was omitted so that the Cia1-Cia2 interaction can be directly interrogated. (**E**) Key to Cia1 variants, colored by substitutions that were nondisruptive (black), partially disruptive (orange) or disruptive (red) of the interaction with Cia2. Also included is a key to the 3 types of negative controls utilized in the copurification experiments.

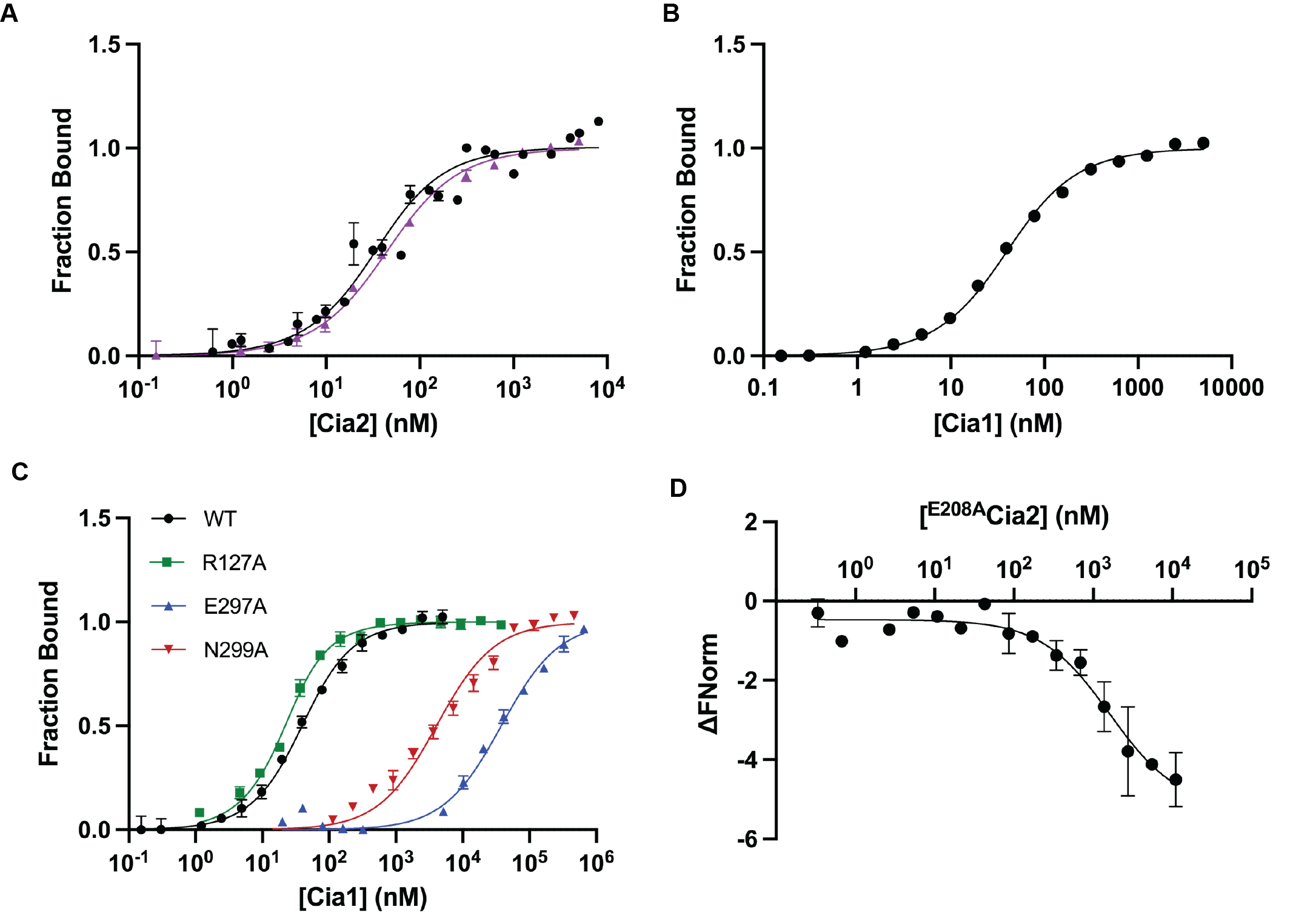

**Figure S3.** Microscale Thermophoresis Assay to monitor *Sc*Cia1-Cia2 complex formation. (**A**) *Sc*Cia1 (20 nM) labeled with RED-NHS was incubated with increasing amounts of either ^DT^Cia2 (black) or ^Δ102^Cia2 (purple). The change in thermophoresis was converted into fraction bound and fit to Eq. 7. ^DT^Cia2 and ^Δ102^Cia2 bind Cia1 with a dissociation constant (*K_D_*) of 24.2±8.41 nM (n=3) and 29.4±0.19 nM (n=3), respectively. (**B**) ^Δ102^Cia2 (20 nM) labeled with RED-MALEIMIDE was incubated with increasing amounts of Cia1 to determine a *K_D_* value of 40±20 (n=3). (**C**) MST assays comparing affinities of the indicated *Sc*Cia1 variants for ^Δ102^Cia2, as in **A**. (D) MST assay of the ^E208A^*Sc*Cia2 variant for *Sc*Cia1, as in **A**.

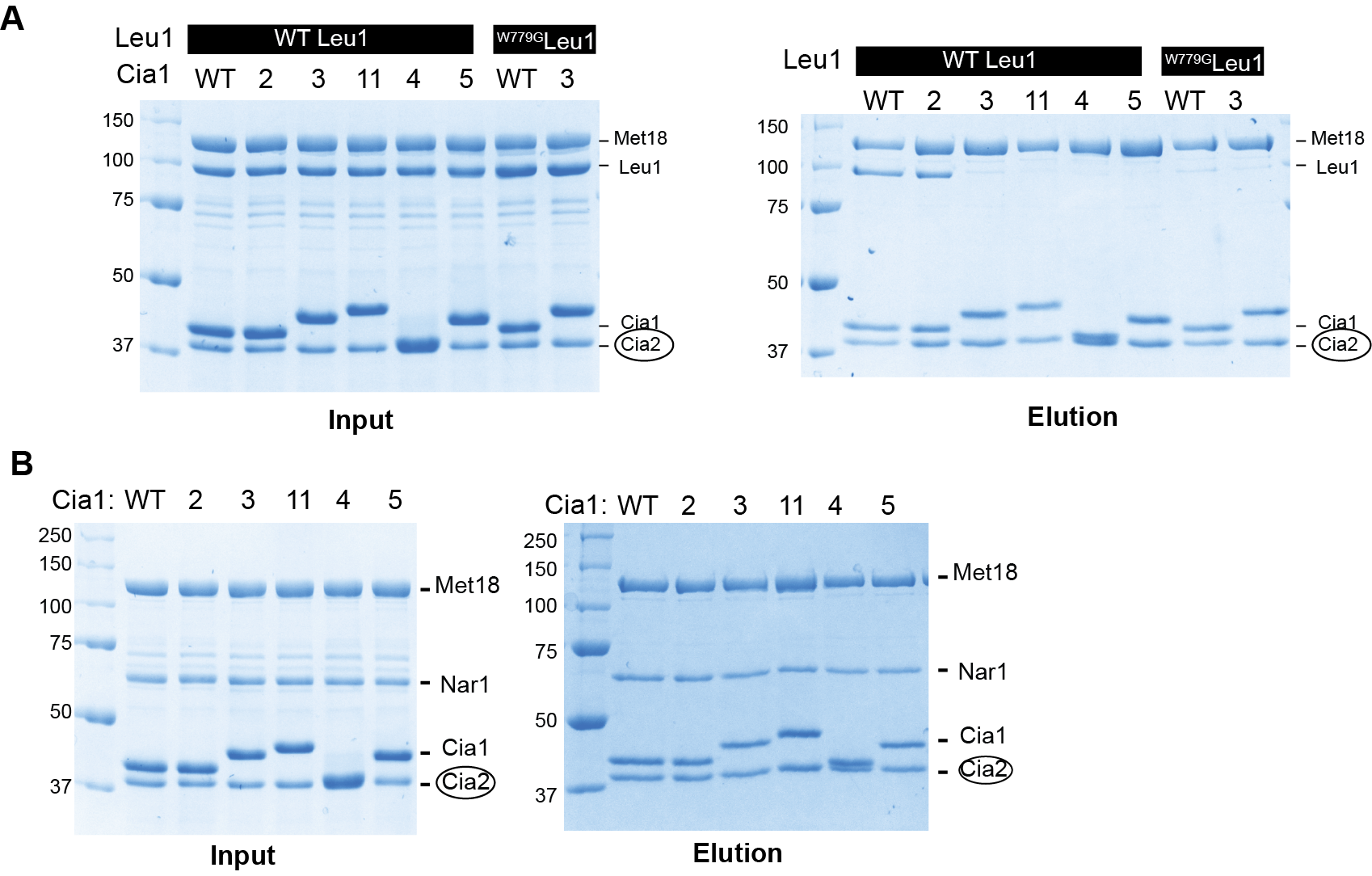

**Figure S4.** Effect of *Sc*Cia1 mutations on *Sc*Leu1 or *Sc*Nar1 binding to the CTC. (**A**) Affinity copurifications in which the strep-tagged *Sc*Cia2 bait (circled) was incubated with *Sc*Leu1 (WT or W779G variant), *Sc*Met18, and the indicated *Sc*Cia1 variant (numbered as in **Figure 2D**). Proteins coeluting from the streptactin resin with the bait were analyzed by SDS-PAGE. The relative migration of Met18, Leu1, Cia1, and Cia2 are indicated to the right of each gel. (**B**) Affinity copurification assays as in **A**, except *Sc*Nar1 was used in place of Leu1.

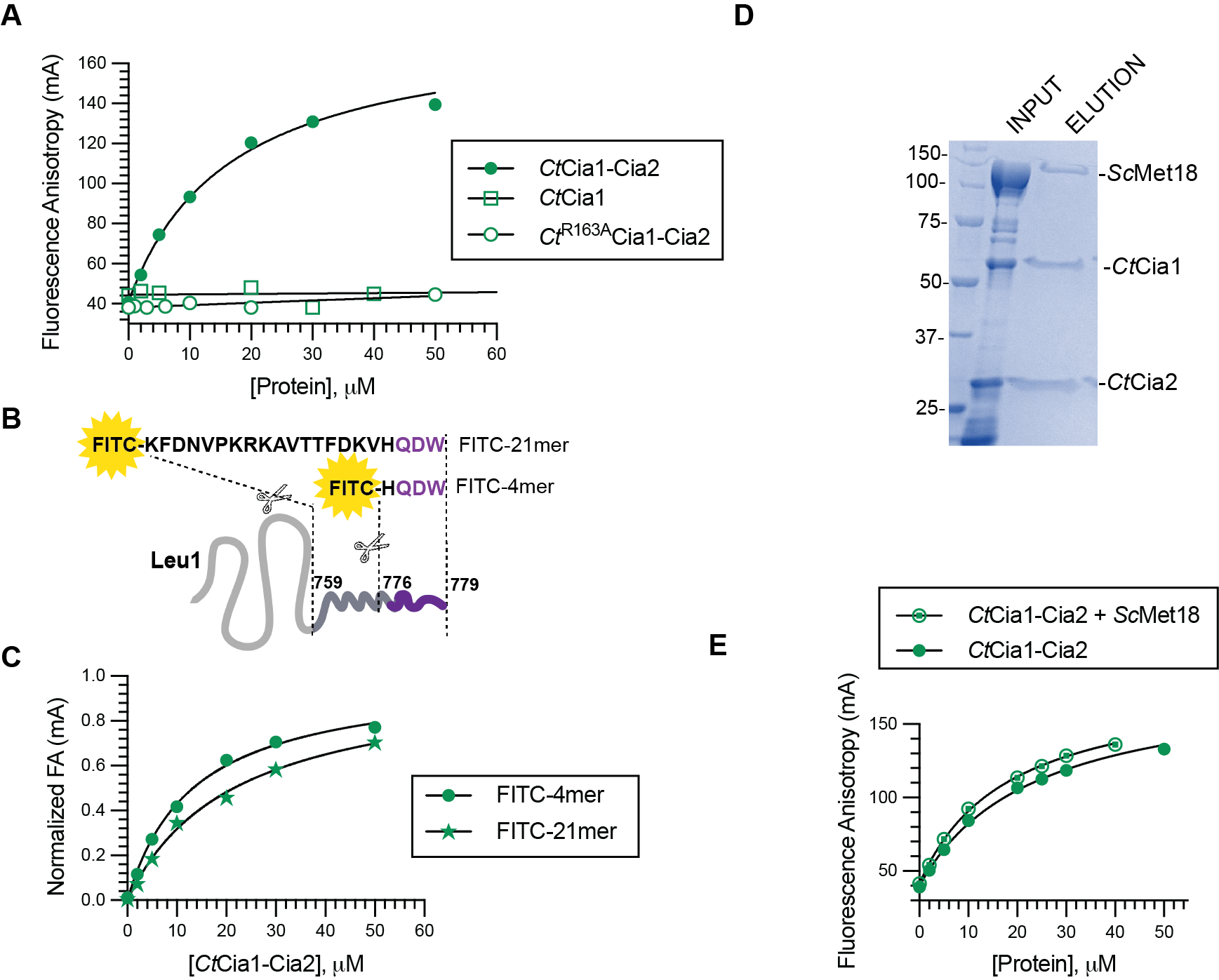

**Figure S5.** FA assay validation for *Ct*Cia1-Cia2. (**A**) FA assays (as in **Figure 3A**) in which the FITC-4mer probe was titrated with the indicated *Ct* protein(s). (**B**) A cartoon representation of the FITC-labeled peptides used in the FA assay. The numbers at the top (759-779) correspond to the *Sc*Leu1’s primary structure. (**C**) FITC-Leu1_4_ or FITC-Leu1_21_ (0.1 µM) were titrated with *Ct*Cia1-Cia2. The normalized FA data (Equation 3) was fit to Equation 2 to determine a *K_D_* value of 13 μM and 21 μM for FITC-Leu1_4_ and FITC-Leu1_21,_ respectively. (**D**) Affinity copurification analysis of *Ct*Cia1-Cia2 and *Sc*Met18 demonstrating that the thermophilic fungal orthologs are able to bind *Sc*Met18. (**E**) The FITC-HQDW (0.1 µM) was titrated with *Ct*Cia1-Cia2 in the presence or absence of an equimolar concentration of *Sc*Met18. The observed apparent *K_D_* values (Equation 2) were 23 µM and 18 µM for the interaction of TCR-peptide with *Ct*Cia1-Cia2 in the absence and presence of *Sc*Met18, respectively.

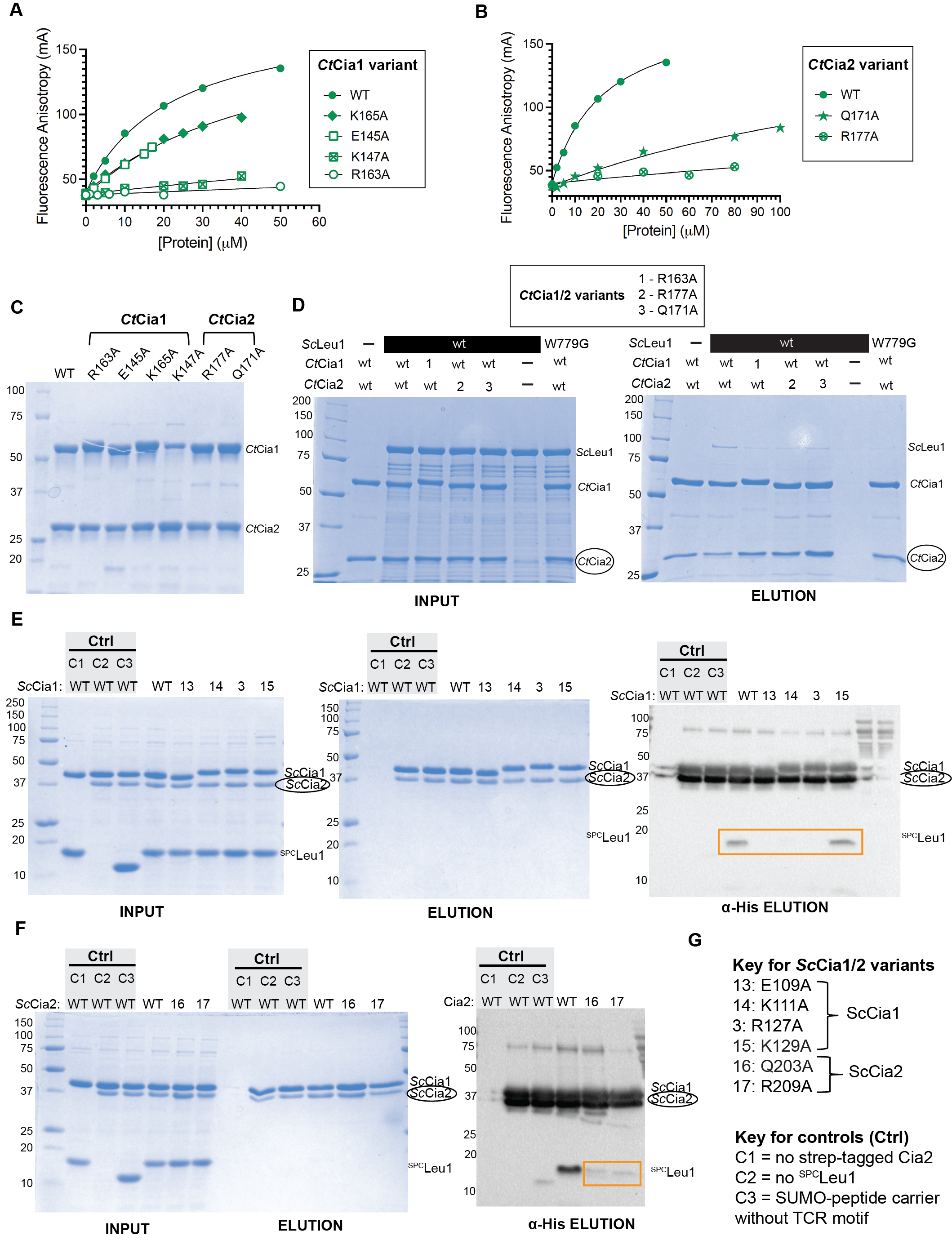

**Figure S6.** TCR peptide interaction with *Ct*Cia1-Cia2 variants. (**A**-**B**) Fluorescence anisotropy binding assays of *Ct*Cia1 (**A**) and *Ct*Cia2 (**B**) variants. Experiments carried out and analyzed as described in **Figure 3A**. (**C**) SDS-PAGE analysis of *Ct*Cia1-Cia2 complex purifications showing that all variants. With the exception of ^K147A^Cia1, all other variants can be purified with similar purity and yield as the WT *Ct*Cia1-Cia2 complex. (**D**) Affinity copurification analysis of *Ct*Cia1-Cia2, or the indicated variant, with *Sc*Leu1 was completed as in **Figure S4A**. (**E**-**G**) Affinity copurification of ^SPC^Leu1 with *Sc*Cia1 (**E**) or *Sc*Cia2 (**F**) variants as indicated in the key (**G**)**.** Regions of western blots shown in **Figure 3E** are boxed.

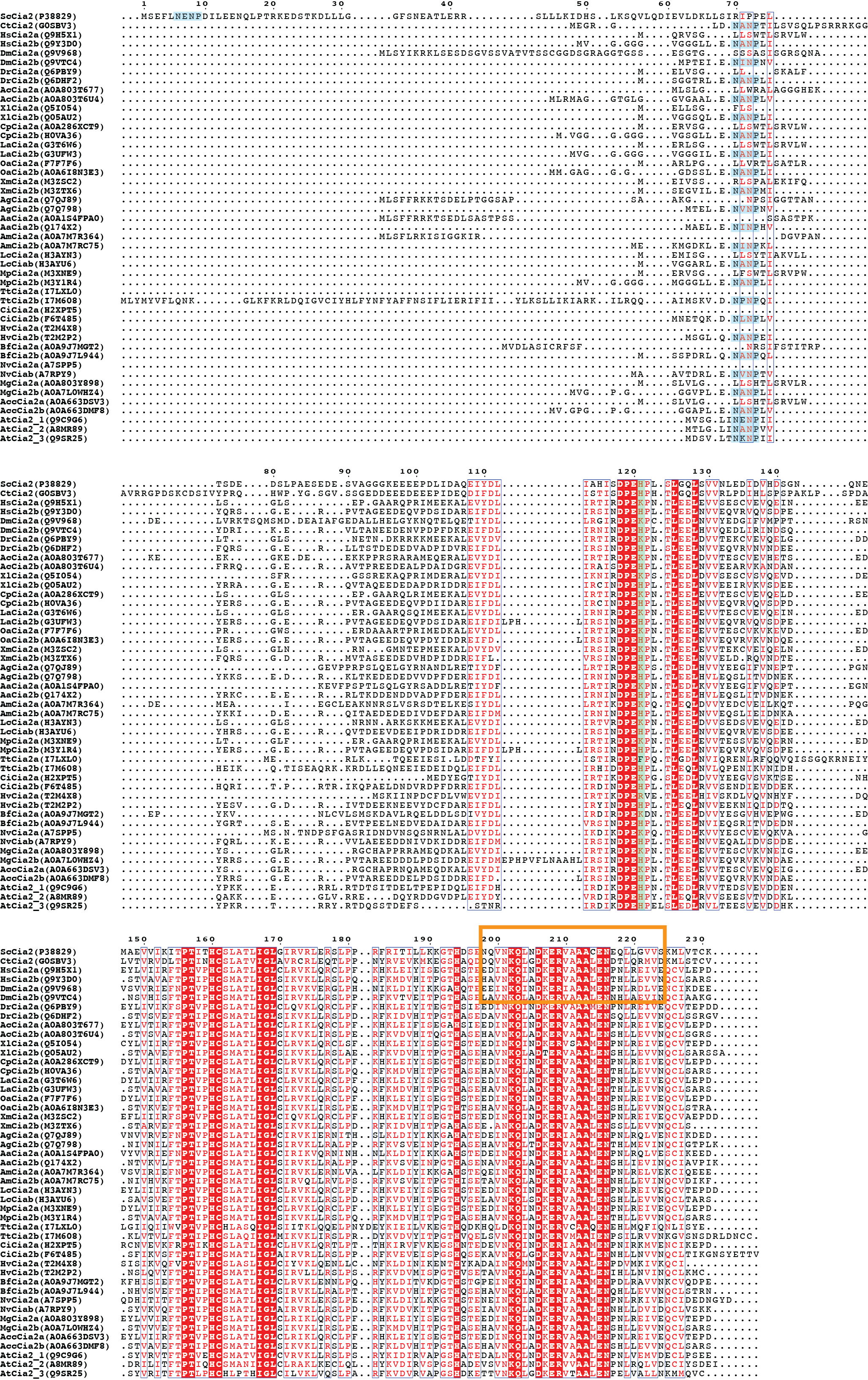

**Figure S7.** Sequence alignment focusing on Cia2a/b pairs. Cia2 sequences from *Ct*, *Sc*, and *At* are compared to 20 Cia2a/b pairs. Uniprot identifies are listed in the label for each sequence. When Cia2a and Cia2b paralogs were not annotated, Cia2b paralogs were identified using the following criteria: the presence of a NxNP motif close to the N-terminus (blue shading), and a His residue immediately following the absolutely conserved DPE motif (green shading). Cia2a paralogs are missing the NxNP motif and they have a Lys residue following the DPE motif. The portion of the alignment shown in **Figure 4A** is boxed in orange, but the entire dataset was used to create the sequence logo shown in Fig 4A.

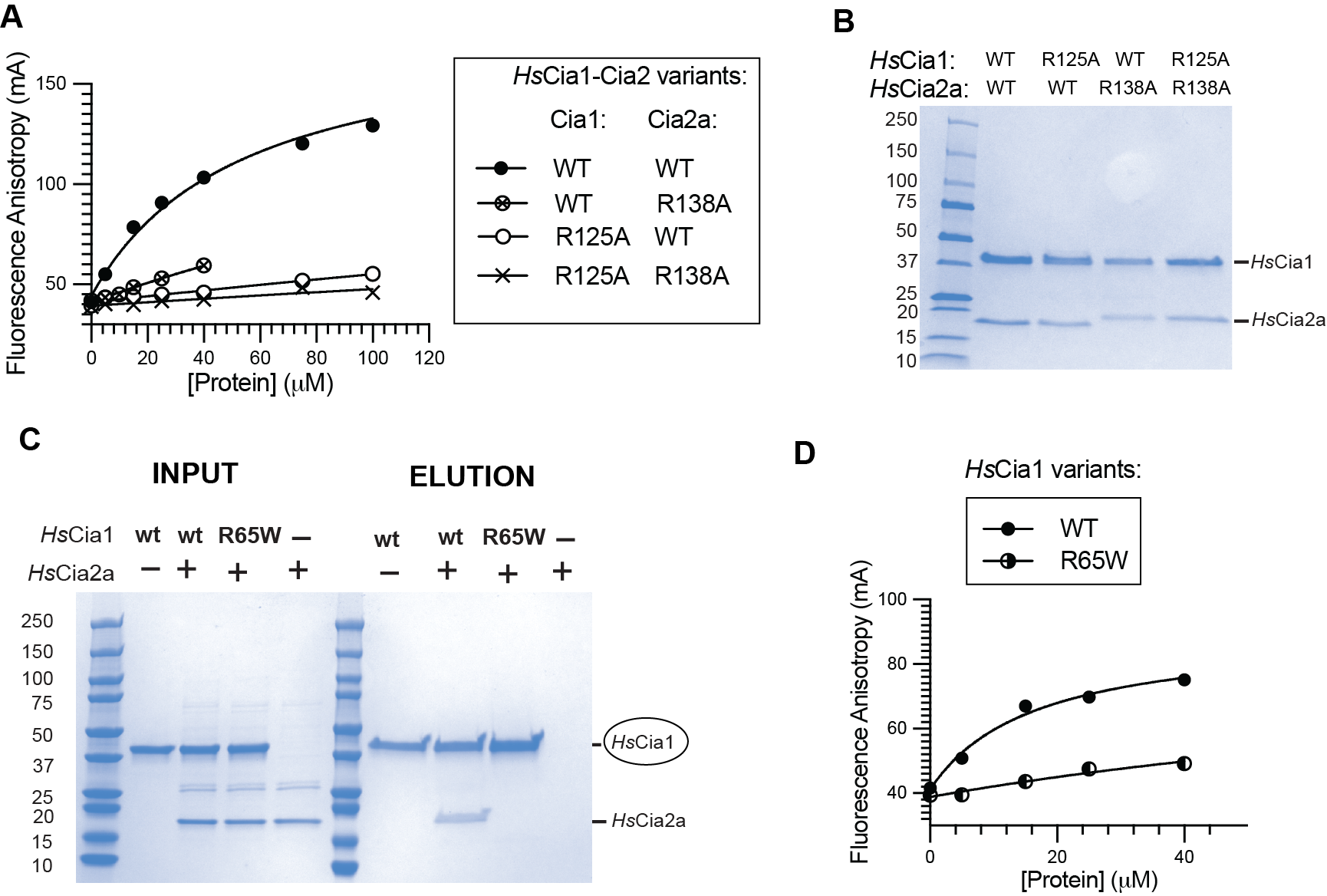

**Figure S8. TCR peptide interaction with *Hs*Cia1-Cia2a.** (**A**) Fluorescence anisotropy binding assays of *Hs*Cia1 and *Hs*Cia2a variants were carried out and analyzed a described in **Figure 4B**. (**B**) SDS-PAGE analysis of *Hs*Cia1-Cia2a complex variants showing none of the Arg variants disrupt formation of the complex. (**C**) Affinity copurification analysis of *Hs*^DT^Cia1 or the R65A variant (bait, circled) with *Hs*Cia2a. (**D**) FA binding assays of *Hs*Cia2a and *Hs*Cia1 variants. *Hs*Cia1 or its R65W variant was titrated into a solution containing 0.1 µM FITC-4mer and 10 µM *Hs*Cia2a. Data plotted and analyzed as in **A** except *A_max_* constrained to 90.
